# Probing the evolutionary dynamics of whole-body regeneration within planarian flatworms

**DOI:** 10.1101/2022.12.19.520916

**Authors:** Miquel Vila-Farré, Andrei Rozanski, Mario Ivanković, James Cleland, Jeremias N. Brand, Felix Thalen, Markus Grohme, Stephanie von Kannen, Alexandra Grosbusch, Han T-K Vu, Carlos E. Prieto, Fernando Carbayo, Bernhard Egger, Christoph Bleidorn, John E. J. Rasko, Jochen C. Rink

**Affiliations:** Department of Tissue Dynamics and Regeneration, Max Planck Institute for Multidisciplinary Sciences, 37077 Göttingen, Germany; Max Planck Institute for Molecular Cell Biology and Genetics, 01307 Dresden, Germany; Georg-August-Universität Göttingen, Johann-Friedrich-Blumenbach Institute for Zoology & Anthropology Animal Evolution and Biodiversity, Germany; Department of Zoology, University of Innsbruck, Innsbruck, Austria; European Molecular Biology Laboratory, Heidelberg, Germany; Department of Zoology & Animal Cell Biology, University of the Basque Country (UPV/EHU), 48080, Bilbao, Spain; Laboratório de Ecologia e Evolução. Escola de Artes, Ciências e Humanidades, Universidade de São Paulo. 02838-000 São Paulo, SP, Brazil; Gene and Stem Cell Therapy Program Centenary Institute, Camperdown, NSW, Australia; Faculty of Medicine and Health, University of Sydney, Sydney, NSW, Australia; Cell & Molecular Therapies, Royal Prince Alfred Hospital, Camperdown, NSW, Australia

## Abstract

Why some animals can regenerate while many others cannot remains a fascinating question. Even amongst planarian flatworms, well-known for their ability to regenerate complete animals from small body fragments, species exist that have restricted regeneration abilities or are entirely regeneration incompetent. Towards the goal of probing the evolutionary dynamics of regeneration, we have assembled a diverse live collection of planarian species from around the world. The combined quantification of species-specific head regeneration abilities and comprehensive transcriptome-based phylogeny reconstructions reveals multiple independent transitions between robust whole-body regeneration and restricted regeneration in the freshwater species. Our demonstration that the *RNAi*-mediated inhibition of canonical Wnt signalling can nevertheless bypass all experimentally tractable head regeneration defects in the current collection indicates that the pathway may represent a hot spot in the evolution of planarian regeneration defects. Combined with our finding that Wnt signalling has multiple roles in the reproductive system of the model species *S. mediterranea*, this raises the possibility of a trade-off between egg-laying and asexual reproduction by fission/regeneration as a driver of regenerative trait evolution. Although initial quantitative comparisons of Wnt signalling levels, reproductive investment, and regenerative abilities across the collection confirm some of the model’s predictions, they also highlight the diversification of molecular mechanisms amongst the divergent planarian lineages. Overall, our study establishes a framework for the mechanistic evolution of regenerative abilities and planarians as model taxon for comparative regeneration research.

## Introduction

Regeneration, defined here as the re-formation of body parts lost to injury, is widespread in the animal kingdom^1^. However, the regeneration abilities of established model systems like *Caenorhabditis elegans, Drosophila melanogaster* or *Mus musculus* are rather limited^2^. In addition, regenerative abilities often vary dramatically between closely related groups or species, e.g., between axolotls (*Ambystoma mexicanum*) and frogs^3,4^, the spiny mouse (*Acomys*) versus the house mouse (*M. musculus*)^5,6^ or within annelids^7^. Between the likely ancestral nature of whole-body regeneration in animals^8^ and the seemingly adaptive effect of regenerative abilities, a fascinating question arises: why is it that only some, but not all, animal species are capable of regenerating lost body parts?

Probing the evolutionary reasons for the variation of regenerative abilities between animals is a challenge because such an attempt necessarily needs to link the molecular and cellular mechanisms that enable regeneration (i.e., proximate mechanisms) to the dimension of ecological or evolutionary mechanisms that select for their presence or absence (i.e. ultimate mechanisms)^9^. As a taxon, planarian flatworms (Platyhelminthes, Tricladida) are especially suitable for a systematic examination of the evolution of regeneration. Many planarian species are capable of whole-body regeneration from tiny tissue fragments. The species *Schmidtea mediterranea (Smed* from now on) and *Dugesia japonica* have been developed into molecularly tractable model species to study the mechanistic underpinnings of whole-body regeneration^10–12^. Regeneration in the model species relies critically on abundant pluripotent stem cells (neoblasts) as the only division-competent cells in planarians^13,14^ and a collectively self-organising network of positional identity signals that orchestrate fate choices^15,16^. The evolutionarily conserved Wnt signalling pathway patterns the planarian A/P axis^17–20^ and the upregulation of Wnt signalling at a wound site is necessary and sufficient for tail specification^19^. Conversely, Wnt inhibition is necessary and sufficient for head specification^18,20,21^. Notably, other planarian species have more limited regenerative abilities or seem to lack a regenerative response altogether^22–24^. Interestingly, the experimental inhibition of canonical Wnt signalling is sufficient for rescuing head regeneration in three planarian species with restricted (regionally-limited) head regeneration^25–27^, indicating that the misregulation of this signalling pathway may contribute to the examined species-specific regeneration defects. With hundreds of planarian species in existence worldwide^28^, a documented range of regenerative abilities from whole-body regeneration to the complete absence of regeneration^23^ and emerging insights into the mechanistic underpinnings of regeneration from the molecularly tractable model species, planarians thus provide a compelling opportunity for investigating the evolution of regeneration.

We here systematically explore the gain and loss of head regeneration across the planarian phylogeny. Central to our approach is a live collection of more than 40 planarian species that we have assembled through systematic field collections and many of which we now maintain in the laboratory. The quantification of head regeneration abilities under controlled conditions in conjunction with a transcriptome-based reconstruction of planarian phylogeny provides a systematic overview of trait evolution within the taxon and uncovers several independent transitions in head regeneration abilities. Our finding that Wnt inhibition rescues even independently evolved head regeneration defects implicates the pathway as a possible trait evolution hot spot. Together with our observation of positive Wnt signalling requirements in the reproductive system of the model species *Smed*, this raises the possibility of a trade-off between egg-laying and asexual reproduction by fission/regeneration as a driver of regenerative trait evolution in planarians. Although comparisons of Wnt signalling levels, regenerative abilities and proxies for investment into sexual reproduction across the collection are consistent with some of the model’s predictions, they also reveal a significant divergence of molecular mechanisms amongst the divergent planarian lineages. Overall, our study highlights the utility of planarians as model taxon for the evolution of regeneration and provides a framework for the analysis of the underlying mechanisms.

## Results

### Establishment of a diverse live collection of planarians

Towards our goal of establishing planarians (Tricladida) as model taxon for studying the evolutionary dynamics of regeneration, we carried out worldwide field collections, preferentially in locations with known high planarian species diversity^29–33^ (Fig. 1a, S. Fig. 1a; see Methods). We mainly focused on freshwater species due to compatibility with the established husbandry protocols of the model species (e.g., *Dugesia japonica* and *Smed*^12,34,35^) and prior reports of species-specific variations in regenerative abilities in the freshwater planarians^23,36^. We developed a combinatorial approach comprising several standardised water formulations, three cultivation temperatures and several different sustenance foods that we routinely try out on field-collected worms (S. Fig. 1b). As a result of this effort, we now maintain a diverse planarian live collection at the MPI-NAT in Göttingen (Fig. 1b-c, S. Fig. 1c-e). Besides the main focus on freshwater planarian species diversity, the collection also represents geographical or morphological population diversity of some species (e.g., multiple isolates of *S. polychroa* from across Europe, including an unpigmented isoline (Fig. 1b)), and some marine species that serendipitously proved amenable to laboratory culture (Fig. 1c). Prolecithophora as a close outgroup to planarians (Fig. 1d), have so far proven difficult to culture in our hands and are not permanently hosted in the collection. Overall, the MPI-NAT collection in its present state represents most major planarian taxa and thus constitutes at least a coarse-grained sampling of extant planarian biodiversity.

**Figure 1.**
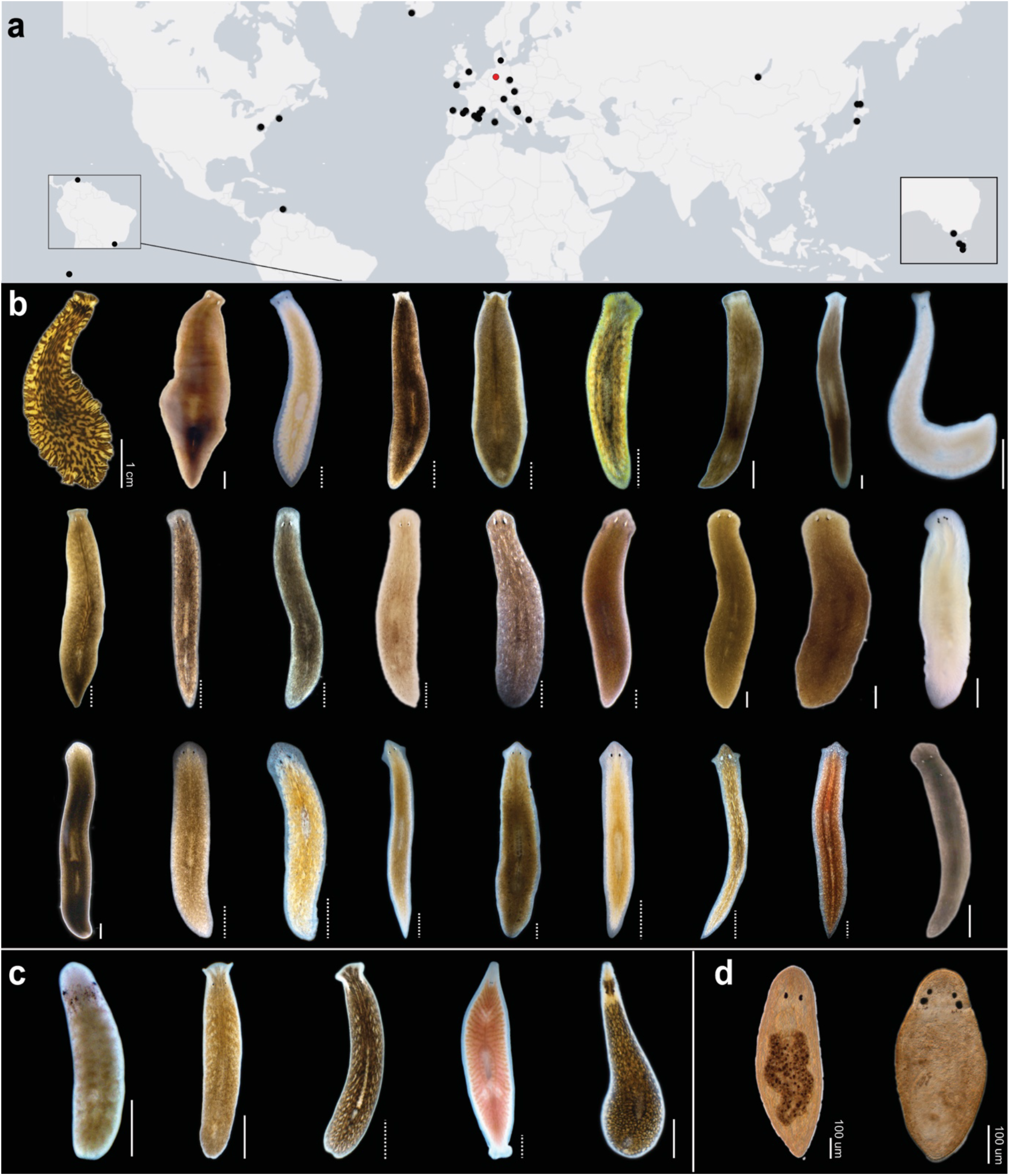
The MPI-NAT planarian collection. **a**, sampling sites (black dots) and collection location (red). **b-d** Live images of flatworms that were characterised as part of this study. **b**, Planarians order Tricladida, Suborder Continenticola. From left to right. First row: *Bdellocephala angarensis; Bdellocephala* cf. *brunnea; Dendrocoelum lacteum; Crenobia alpina; Polycelis felina; Polycelis tenuis; Polycelis nigra; Seidlia* sp.; cf. *Atrioplanaria*. Second row: *Phagocata gracilis; Hymanella retenuova; Phagocata pyrenaica; Planaria torva; Cura pinguis; Cura foremanii; Schmidtea lugubris; Schmidtea polychroa* (dark strain); *Schmidtea polychroa* (unpigmented strain). Third row: *Schmidtea nova; Smed* (asexual strain); *Dugesia tahitiensis; Dugesia sicula; Dugesia* sp.; *Dugesia japonica; Girardia dorotocephala; Girardia tigrina; Spathula* sp.3. **c**, *Planarians*, Suborder Maricola. *Camerata robusta; Procerodes littoralis; Procerodes plebeius; Bdelloura candida; Cercyra hastata*. **d**, Order Prolecithophora: *Plagiostomum girardi; Cylindrostoma* sp. Scale bar: 1 mm unless otherwise noted; measured (pixel resolution; solid line) or approximated during live imaging (graph paper; dotted line).

We next took advantage of our species collection to systematically compare regeneration abilities under standardised laboratory conditions. We focused on head regeneration as the arguably most complex planarian regeneration challenge (e.g., brain re-formation and rewiring), for which the regeneration of the eye spots provides a convenient morphological readout. Specifically, for each species, we assayed the fraction of successful head regeneration at five transverse cuts equally spaced along the A/P axis, inspired by Šivickis’ pioneering studies^23^, (Fig. 2a; see Methods). Regeneration defects were highly reproducible for a given species and generally consistent with previous literature reports. Our screen identified three broad groups of head-regeneration capacities (Fig. 2a, b). The first group, designated in the following as “robust regeneration” (= group A), comprises species with high head regeneration success all along the A/P axis (≥ 80% surviving fragments, irrespective of cut position). The head regeneration frequency plots along the body axis for these species, termed “head frequency curve” in the planarian literature^24^, are flat and uniformly high (Fig. 2b). Two prominent members of this group are the two planarian model species *Smed* (both the fissiparous and the egg-laying strain (Fig. 2a, S. Fig. 2a)) and *D. japonica*, which were originally selected based on their robust and rapid regeneration. Despite their robust head regeneration all along the A/P axis, group A species may still display A/P differences in the rate of head regeneration, for example, in *Smed*^37^. The second group, designated here as “restricted regeneration” (= group B), comprises species with positional head regeneration defects, which we define here as <80% head regeneration efficiency at one or more A/P axis positions (Fig. 2a, b). This group includes freshwater species with known deficiencies in head regeneration in the posterior body half (e.g., *Dendrocoelum lacteum*)^23^, several so far undescribed regeneration defects (e.g., *Cura pinguis*), and also some marine species (e.g., *Cercyra hastat*a). The “restricted regeneration” group also includes several cases of reduced regeneration efficiency in central regions, e.g., in *Dugesia* sp. (Fig. 2a, b). The third group, designated in the following as “poor regeneration” (= group C), comprises species that are incapable of head regeneration in any amputation fragment under our experimental conditions, with the marine planaria *Bdeloura candida* as a known example (Fig. 2a, b)^23^.

**Figure 2.**
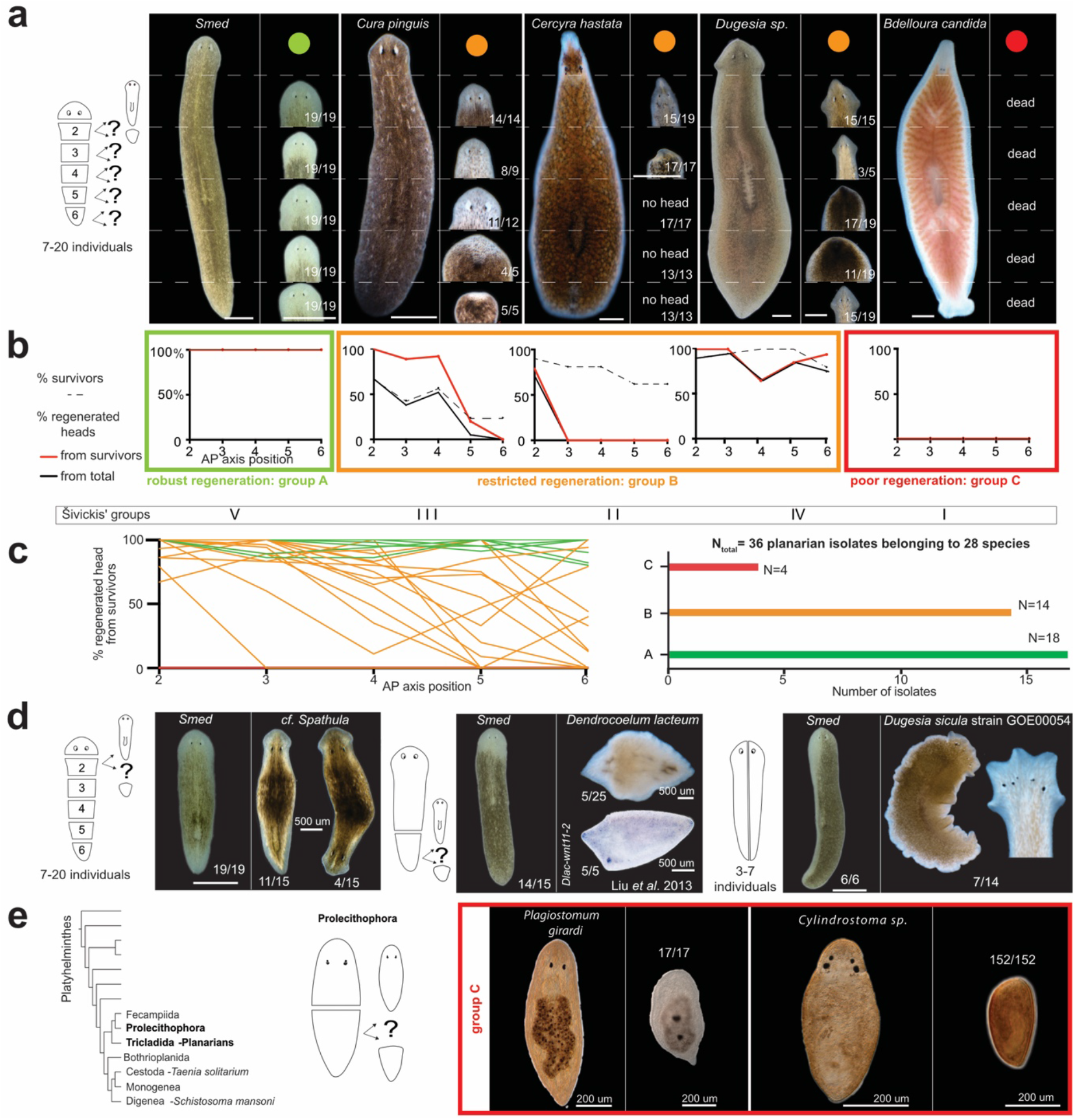
Quantitative analysis of head regeneration abilities across the flatworm collection. **a**, Cartoon of the serial head regeneration assay (left) and representative outcomes in the indicated species. Dashed lines: amputation planes and typical regeneration outcomes (small images) and their relative frequency (number pairs; “dead” in case of no survivors) at the respective A/P position. Colors designate the head regeneration classification scheme used throughout this study, e.g., group A/green: “robust regeneration” (efficient head regeneration at all A-P axis positions); Group B/orange: “restricted regeneration” (position-dependent head regeneration defects); Group C/red: “poor regeneration” (no head regeneration at any A/P position). Missing images in the *C. hastata* panel correspond to pieces that did not regenerate a head and died shortly before imaging. **b**. Graphical representation of the data in the form of head frequency curves. The percentage of successful head regeneration is plotted either as a fraction of the initial (black line) or only surviving fragments (red line). Dashed line: percentage of fragment survival. The roman numerals below designate the more fine-grained head-regeneration scheme by Šivickis^90^. **c**, Left: Head frequency curves (percentage of surviving fragments) for all analysed species and color-coded as in a; right: number of species in each head regeneration category. **d**, Unclassified regeneration defects and their relative frequencies in the indicated collection species, including bi-polar double-heads (left), double-tails (centre) and mediolateral axis duplication (right). Amputation paradigms are cartooned to the left; the respective regeneration outcome in *Smed* is shown for reference. **e**, Left: Phylogenetic overview of flatworm groups related to planarians^40^. Centre: Amputation paradigm and right: representative images and quantifications of head regeneration failures and in the indicated Prolecitophora species (right). Scale bar: 1 mm unless otherwise noted.

Out of the 36 analysed planarian field isolates, only four were classified as poor regenerators (all different marine species), 14 were assigned to the “restricted regeneration” class, and a total of 18 isolates displayed “robust regeneration” (Fig. 2c). Significantly, our collection also harbours examples of each class in Šivickis’ more fine-grained regeneration classification scheme^23^, thus providing the intended broad overview of planarian regeneration and regeneration defects (Fig. 2b). Further, the collection also holds multiple examples of interesting species-specific regeneration peculiarities, such as a propensity for bipolar head or tail regenerates in *Spathula* sp. and *Dendrocoelum lacteum* or frequent aberrant axis duplications in mediolateral regenerates of one population of *Dugesia sicula* (Fig. 2d). Interestingly, we also found the model species *Dugesia japonica* to regenerate double-headed individuals with low frequency (S. Fig. 2b), which is relevant regarding the interpretation of previously reported regeneration experiments in space^38,39^. Although not captured by our classification scheme, each of these species-specific regeneration peculiarities represents a future opportunity for probing the mechanistic underpinnings of planarian regeneration.

To put the regenerative abilities of planarians into a phylogenetic perspective, we further included Fecampiidae and Prolecitophora into our analysis, which together form the sister taxon to Tricladida^40,41^. While we could neither obtain live specimens nor published information on the regenerative abilities of the Fecampiidae, we obtained live specimens of two Prolecithophora species (Fig. 2e). Due to their small size (<1 mm length), we could only examine head regeneration upon medial bisection. None of the posterior pieces regenerated a new head under our assay conditions. This result is in agreement with a recently published analysis^42^ and confirms that Prolicetophora are iapable of *de novo* head regeneration, but only can repair injuries to an existing head similar to *Macrostomum* or polyclads^43,44^. The rich and varied regeneration of planarians, therefore, contrasts with so far poor regeneration abilities of the sister groups, thus making the evolutionary history of regeneration in planarians particularly interesting^34^.

An essential prerequisite for assessing trait evolution is a comprehensive understanding of phylogeny. Although multiple phylogenies of the Tricladida have been published to date, they are either based on morphological evidence or cover a limited number of species or gene sequences^28^. Towards complementing these data, we carried out short-read RNAseq and de-novo transcriptome assembly of 41 planarians and sister group species using our published pipeline (Fig. 3a)^45,46^. Additionally, we incorporated publicly available or contributed data from eight species, yielding a total data set of 49 transcriptomes (45 planarians and four outgroup Platyhelminthes). To quality control our transcriptomes, we carried out a BUSCO (Benchmarking Universal Single-Copy Orthologs)^47^ analysis, using all available flatworm and selected outgroup transcriptomes in ENSEMBL^48^ as a reference. The consistently high completeness and low fragmentation of BUSCO gene copies in our transcriptomes indicated a high assembly quality of our data set (Fig. 3b). However, the total number of BUSCO genes in flatworm transcriptomes was consistently lower than in outgroup species. Interestingly, this discrepancy can be attributed to 109 genes absent from 90% of the total flatworm transcriptomes (Fig. 3c). Of those, 74 genes were present in the most distant outgroups to planarians in our analysis, *Macrostomum lignano* or *Prostheceraeus vittatus* or both, yet largely absent in planarian and close outgroup transcriptomes, prolecitophora and fecampiidae (Fig. 3c, inset for examples; S. Table. 1). These 74 BUSCO genes, therefore, represent likely gene losses in planarians and related flatworm groups, thus further corroborating the previously reported substantial gene loss in the genome of the model species *Smed*^49^. Altogether, our analysis highlights both, the need for better integration of so far undersampled taxonomic groups into BUSCO and the high assembly quality of our flatworm transcriptomes, which will be deposited and made publicly available in the online repository Planmine^45^.

**Figure 3.**
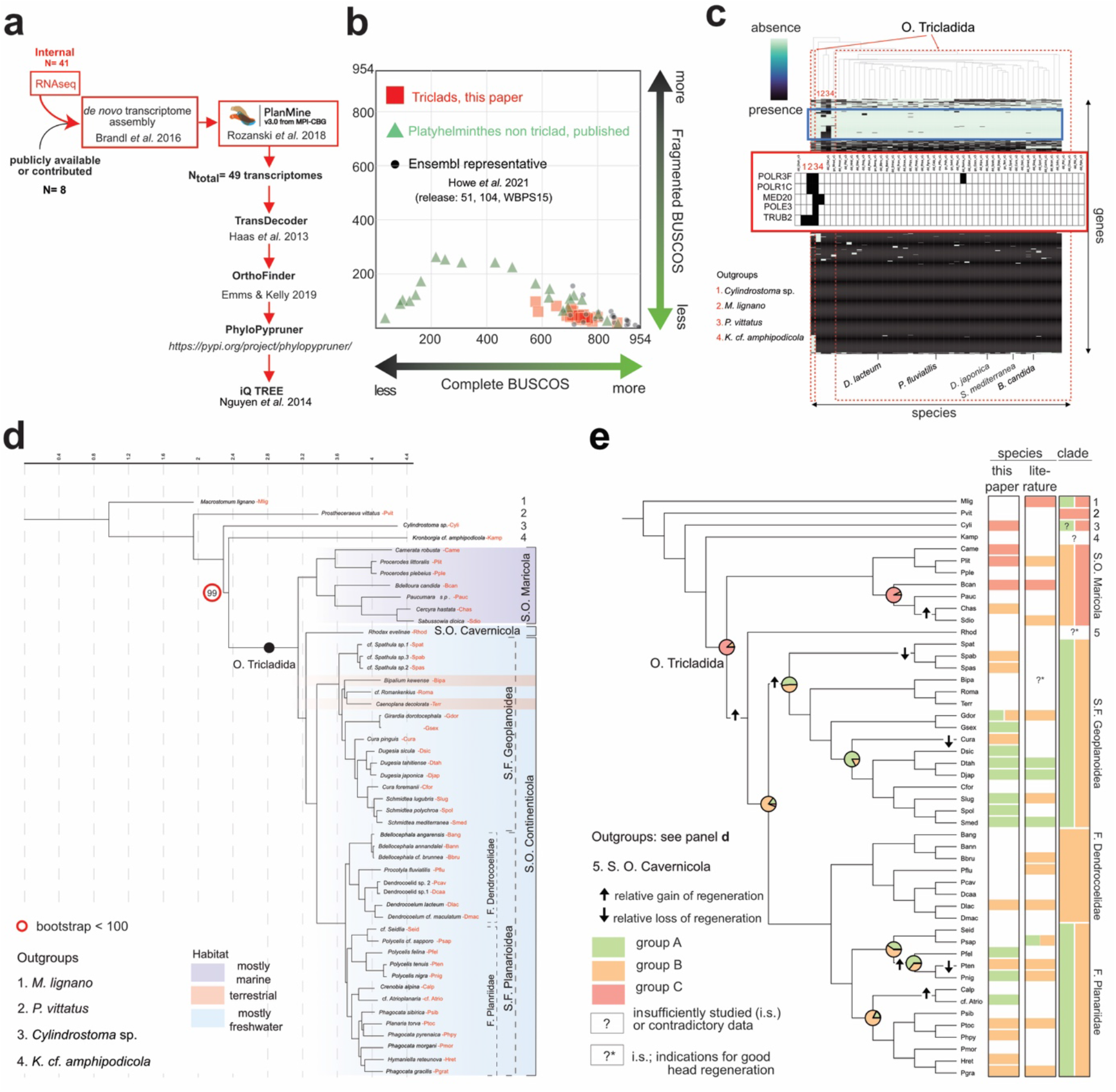
Phylogenetic analysis of head-regeneration abilities in planaria. **a**, *de novo* high-quality transcriptome assembly pipeline and unbiassed gene sequence selection workflow for multi-gene phylogeny reconstruction. **b**, BUSCO quality comparison between the indicated transcriptome sources, plotting BUSCO gene sequence completeness (X-axis) versus fragmentation (y-axis). “Perfect” BUSCO representation: Bottom right corner. **c**, Presence (black)/absence (turquoise) analysis of individual BUSCO genes (Y-axis) in flatworm transcriptomes (X-axis, in phylogenetic order) and representative outgroups. Names of representative species are indicated (see S. Table 1 for detail). The red frame designates BUSCO genes absent from > 90% of the transcriptomes. The blue inset shows a selection of likely planarian-specific BUSCO loss events. **d**, Maximum-Likelihood (ML) tree on basis of the transcriptomes with representatives of all the suborders of the Tricladida. Numbers indicate outgroups, and the preferred habitats of major planarian taxonomic groups are indicated by color shading (see legend). **e**, Phylogenetic map of planarian head regeneration abilities and ancestral state reconstruction (ASR). Pie charts at nodes represent the proportion of character histories with the indicated state for regenerating capability (see S. Fig. 3c, S. Table 2 and 3, and methods). The color coding of the columns to the right indicates species- and clade-specific head regeneration abilities quantified by this study or extracted from the literature (both used for clade annotation).

To comprehensively reconstruct the phylogeny of our planarian species collection, we used the pipeline cartooned in Fig. 3a to extract broadly conserved single-copy orthologues from our transcriptomes^50–52^. The resulting dataset of 79312 sequences organised into 1684 alignments across 49 species includes so far poorly investigated planarian genera (e.g., *Cura, Camerata* or *Hymanella*) and species (e.g., *Phagocata morgani*) and represents the most extensive taxon sampling to date. Accordingly, the Maximum-Likelihood representation of the phylogenetic tree shows very high support for all branches (Fig. 3d). A coalescent-based species tree estimation strategy recovered a similar tree topology (Astral; S. Fig. 3a). Our phylogeny is broadly consistent with the previously inferred relations between major planarian clades^28^. One interesting exception is the placement of the taxon Cavernicola (Fig. 3d), which recent molecular analysis considered a sister group of the Maricola^53^. In contrast, our phylogeny indicates a sister group position to the Continenticola. Although based only on a single species representative, the high number of genes analysed and the high bootstrap value (100) support this new phylogenetic hypothesis, which deserves further investigation. Further, noteworthy revisions include, for example, the inferred split of the two *Cura* species, which do not form a monophyletic group, or the relation between terrestrial planarians and the freshwater genus cf. *Romankenkius*^54^. While a detailed taxonomic analysis will be published elsewhere, a striking feature of our phylogenetic tree is the deep splits between different planarian lineages. Branch length comparisons with selected mammalian and nematode transcriptomes quantitatively confirmed the extreme divergence of planarian lineages, which exceeds the observed sequence divergence between mammalian species and is on par with that of nematodes (S. Fig. 3b). Whether the high sequence divergence reflects the old age of planarian lineages or unusually rapid rates of genome evolution is difficult to ascertain due to the poor fossil record of the group. Overall, our phylogeny establishes a resource for systematic and phylogeographic studies and the necessary framework for comprehensive analyses of planarian trait evolution.

To reconstruct the evolutionary history of planarian head regeneration, we combined the phylogeny and the above quantitative analysis of head regeneration abilities (Fig. 3e, S. Fig. 3c and further literature mining^23,36,55–57^) for an ancestral state reconstruction (ASR). At least two other flatworm clades (Catenulida and Macrostomorpha) harbour species capable of *de novo* head regeneration (S. Fig. 3d)^44^. Both are distantly related to planarians, and the information available on the regenerative abilities of more closely related sister groups is limited (see above). In this context, ASR best supports the following trait evolution scenario: the complete absence of head regeneration (“poor regeneration”; red) as the most likely ancestral state in planarians, followed by the acquisition of regionally limited head regeneration (“restricted regeneration”; orange) in the Continenticola and isolated marine lineages and the final acquisition of robust position-independent head regeneration (“robust regeneration”; green) exclusively in freshwater lineages. Transitions between poor regeneration and restricted regeneration are rare in our data set (2) and only observed in the marine species (Maricola) and at the base of the clade, grouping the rest of the planarian species (Fig. 3e and S. Tables 2 and 3). On the other hand, transitions between “restricted regeneration” and “robust regeneration” appear to be limited to the Continenticola, but likely occur frequently and in both directions (particularly in the Planariidae; Fig. 3e and S. Table 3). Importantly, our present taxon sampling cannot address whether *de novo* head regeneration is ancestral in the Platyhelminthes. More extensive taxon sampling of the different flatworm taxa and, ultimately, also mechanistic comparisons of the head regeneration process in Catenulids, Macrostomorppha and Planarians will be required. However, for planarians, our data demonstrate an unexpectedly dynamic evolutionary history of head regeneration capabilities, with at least five independent gains and three independent reductions of head regeneration capability supported by our present ~40-species survey (Fig. 3e and S. Table 3).

Independent evolutionary transitions in head regeneration abilities imply independently evolved changes in the underlying molecular control network. So far, misregulation in the highly conserved Wnt signalling pathway has been linked to the head regeneration defects of three planarian species (two dendrocoelids *D. lacteum and Procotyla fluviatilis* and the planariid *Phagocata kawakatsui*)^25–27^. Our live species collection provided a unique opportunity to systematically examine the contribution of Wnt signalling to the many independently evolved head regeneration deficiencies in planarians. As we have previously shown in *Smed*, planarian canonical Wnt signalling activity can be monitored via the quantification of *ß*-CATENIN-1 amounts due to the functional segregation of the ancestral signalling and cell adhesion roles of *ß*-CATENIN between different gene homologues^58^. Western blotting with our previously characterised anti-*Smed*-*ß*-CATENIN-1 antibody raised against the highly conserved armadillo repeat region of *Smed*-*ß*-CATENIN-1^17^ detected a band of the expected size (~120 kDa) in multiple collection species (Fig. 4a, S. Fig. 4c), thus providing a likely proxy for pathway activity beyond the model species. To modulate canonical Wnt pathway activity in the different species, we targeted the respective *ß-Catenin-1* (S. Fig. 4a, b) and *Adenoma Polyposis Coli (APC; a* core component of the ß-CATENIN destruction complex) homologues by feeding with in vitro translated dsRNA, which in the model species *Smed* results in a decrease or increase of ß-CATENIN-1^17^ levels (Fig. 4b). *C. pinguis, Planaria torva, C. robusta* and *B. candida* displayed broadly similar changes in relative ß-CATENIN-1 levels in response to the *RNAi* treatments, thus indicating the principal utility of the *RNAi* approach for experimental alterations of Wnt pathway activity in those species. Interestingly, the species *P. tenuis* was largely refractory to *RNAi* by feeding (Fig. 4b) and even injection (not shown). This observation provides a first experimental indication that not all planarian species may be equally susceptible to systemic *RNAi*, similar to the previously observed species-specificity of the *RNAi* response in nematodes^59,60^.

**Figure 4.**
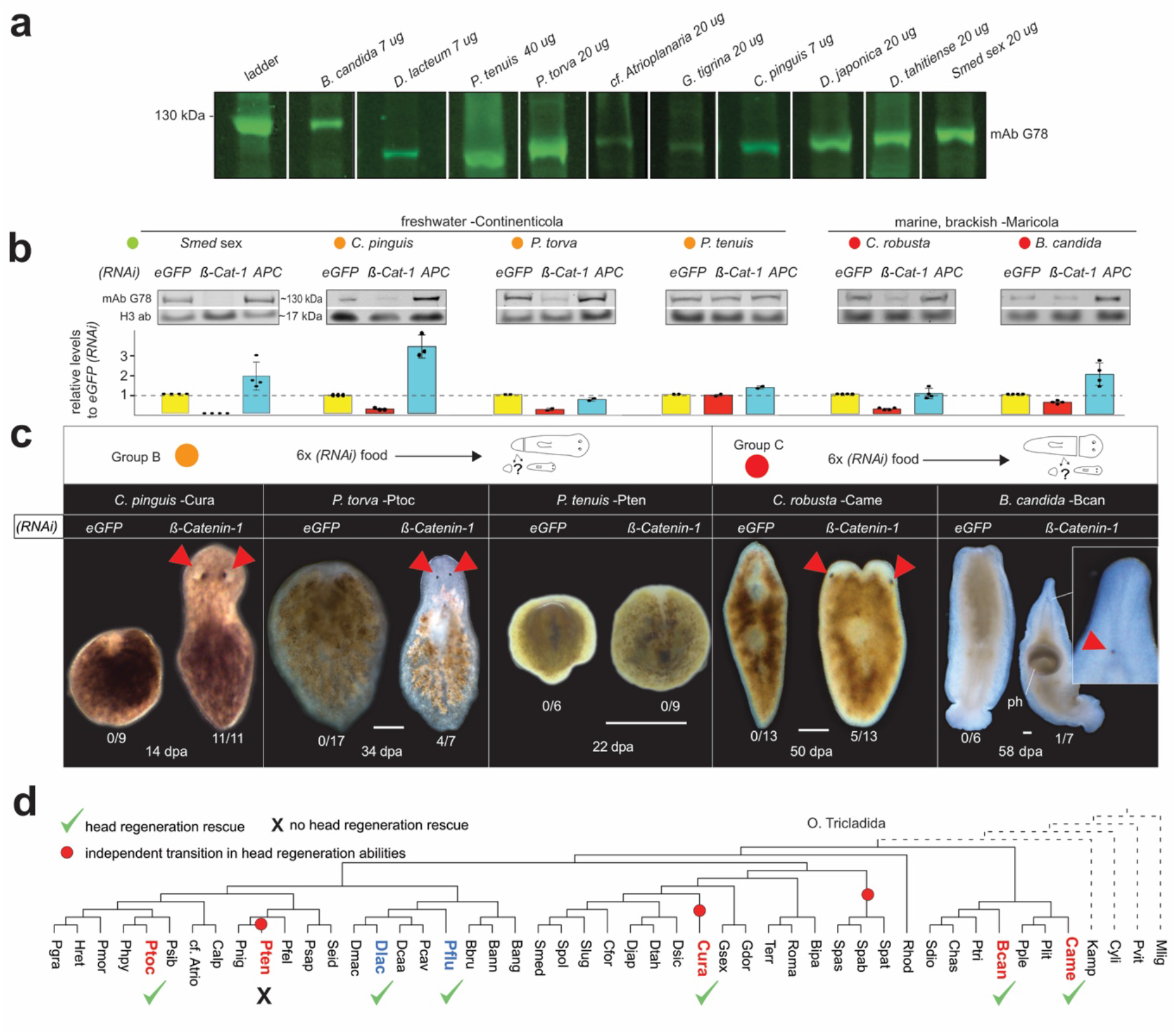
Canonical Wnt pathway inhibition rescues head regeneration defects across planarian phylogeny. **a** Fluorometric Western blot demonstration of anti-*ß*-CATENIN-1 antibody G78 crossreactivity with tail tip lysates of different planarian species. Species and amount of lysate/lane as indicated. **b**, RNAi-mediated Wnt pathway activity modulation in the indicated species from regeneration groups A, B or C (color coding). Animal cohorts were treated with *eGFP(RNAi) (*control), *APC(RNAi)* (gain of Wnt signalling), or *ß-Catenin-1(RNAi)* (loss of Wnt signalling; see “Methods”). Top: Representative quantitative Western blots for *ß*-CATENIN-1 and Histone H3 as loading control. Bottom: Bar graph representation of G78 signal intensity relative to the control in each species (*eGFP(RNAi)*). Error bars denote the SD of four technical replicates (dots) of a representative experiment. **c**, head regeneration rescue assay upon Wnt inhibition/*ß-Catenin-1(RNAi)* in the indicated category B or C species. Live images at the indicated day post-amputation (dpa) and *RNAi*-conditions; number pairs indicate the observed frequency of head or eye regeneration. Red triangles: regenerated eyes. Scale bar: 500 um. **d**, Phylogenetic representation of documented head regeneration rescued by *ß-Catenin-1(RNAi)* (green check-mark) (phylogeny extracted from Fig. 3d). This study (red): *C. pinguis, P. torva, C. robusta* and *B. candida*. Previous studies (blue): *D. lacteum*^25^ and *P. fluviatilis^26^. Phagocata kawakatsui*, for which head regeneration rescue was also reported^27^, was excluded due to the lack of a publicly available transcriptome.

To assess the Wnt signalling dependence of head regeneration defects, we next assayed the sufficiency of *ß-Catenin-1(RNAi)* to rescue head regeneration. With the dugesiid *C. pinguis*, the planariids *P. torva* and *P. tenuis* as members of the restricted regeneration group, and the maricola *C. robusta and B. candida* in the poor regeneration group, our species set comprised representatives of the major phylogenetic branches and multiple independently evolved transitions in regenerative abilities (see Fig. 3e). *ß-Catenin-1(RNAi)* indeed rescued head regeneration or promoted the appearance of head-like structures in all species except the *RNAi*-response deficient *P. tenuis* (see above), yet with varying efficiencies (e.g., 11/11 in *Cura pinguis* and 1/7 in *B. candida*). Even the lower rescue efficiencies were relevant due to the complete absence of similar phenotypes in the corresponding control pieces and, therefore, likely reflect the varying knock-down efficiencies in the different species (Fig. 4c). Intriguingly, we even observed indications of head-like tissue formation upon *ß-Catenin-1(RNAi)* in the two maricola species tested, which are likely ancestral group C members (poor regeneration) and maximally distant to the model species *Smed*. When considered in the context of planarian phylogeny and published data, our results indicate that the inhibition of canonical Wnt signalling via *ß-Catenin-1(RNAi)* can generally bypass head-regeneration defects in planarians, independent of the evolutionary history of the specific lineage (Fig. 4d). By extension, these results further imply that head regeneration defects across planarian phylogeny may be associated with a functional excess of Wnt pathway activity, thus making this pathway a putative hot spot in the evolution of planarian head regeneration defects.

Regeneration-deficient planarian species are not laboratory mutants. Therefore, natural selection scenarios must exist in which regeneration deficiencies become selectively neutral or even positive. A common hypothesis is that robust whole-body regeneration abilities are under selection as a necessary aspect of asexual reproduction by fission^44,61^, which is also one of the two reproduction modes available to planarians. As exemplified by the obligate fissiparous strain of the model species *Smed* (commonly referred to as the asexual strain; Fig. 5a right), many planarians reproduce by splitting themselves (fission), followed by whole-body regeneration of the two pieces. Alternatively, planarians can also reproduce via the deposition of large yolk-rich egg capsules, as exemplified by the egg-laying strain of the model species *Smed* (commonly referred to as the sexual strain; Fig. 5a left). Reproduction via egg-laying can occur independently of the regenerative abilities of the species, but species with restricted or poor regeneration always replicate via egg-laying (e.g., the Maricola) (S. Fig. 5a). To further explore the putative link between regeneration and reproduction in planarians, we examined the taxon and habitat distribution of reproduction modes and regenerative abilities in the literature and our field notes. As shown in Fig. 5b, egg-laying and poor or restricted regeneration tend to associate at the taxon level, while fissiparous strains or species are almost invariably robust regenerators (Fig. 5b; the mild central body regeneration restriction in some dugesids (e.g., *Girardia dorotocephala*) is the only exception). Furthermore, poor or restricted regeneration species tend to occur in stable habitats (e.g., larger rivers, lakes or the sea), while robustly regenerating fissparous strains or species tend to occur in less stable habitats (e.g., fast-flowing streams or temporary water bodies (S. Fig. 5a^62,63^)). Hence our data are consistent with the hypothesised selection of robust whole-body regeneration as a necessary physiological correlate of reproduction by fission, which may thus become dispensible in habitats that favour egg-laying species.

**Figure 5.**
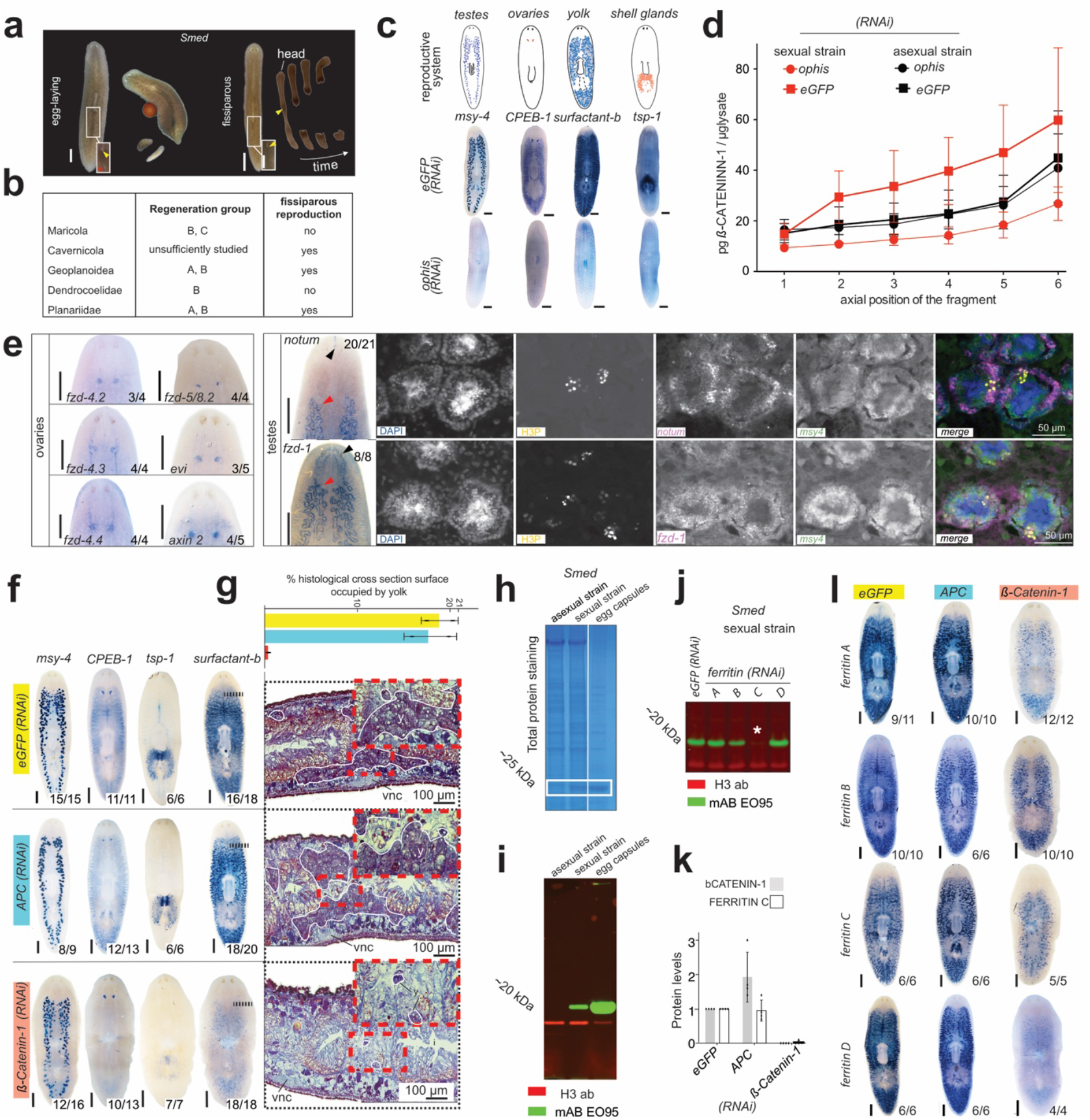
Diverse roles of Wnt signalling in the reproductive system (RS) of *Smed*. **a**, Live image montages illustrating the reproduction modes of the two *Smed* laboratory strains, egg-laying in the sexual strain (left) and fission/regeneration in the asexual strain (right). Insets: Presence/absence of the ventral gonopore (red star) posterior to the moth opening (yellow arrowhead) as a distinguishing feature. **b**, Correlation between robust head-regeneration/Group A and fissiparous reproduction across planarian clades. Data synthesis based on literature, Rink lab field notes, and MPI-NAT live collection observations. **c**, Cartoon representation (top) and colorimetric whole-mount in situ hybridisations with the indicated probes (middle) of the major elements of the hermaphroditic planarian RS; bottom: Loss of all RS elements upon *ophis(RNAi)*. **d**, Wnt signaling activity/*ß*-CATENIN-1 abundance in six serial A/P sections (1 = head; 6 = tail tip) in the indicated *Smed* strain and RNAi condition. *ß*-CATENIN-1 levels were quantified by quantitative Western blotting with the G78 mAB and normalized to the Histone H3 signal in each lane and an external standard. Error bars denote the SD of four biological replicates, each represented by the mean of three to four technical replicates. **e**, Expression patterns of the indicated Wnt pathway components in the *Smed* reproductive system by colorimetric or fluorescent wholemount in situ hybridisation. Number pairs represent the observation frequency of the pattern shown; red arrowhead: testes; black arrowhead: labelling of elements out of the RS; H3P: Antibody staining of the M-phase marker Histone H3 Ser10 phosphorylation. **f**, Colorimetric whole-mount in situ hybridisation expression patterns of the indicated reproductive system marker probes under control (*eGFP), APC* or *ß-Catenin-1(RNAi)*. Number pairs: Observation frequencies of the pattern shown. Dashed lines: approximate position of the sagittal section in Fig. 5g. **g**, quantification of the relative cross-sectional area occupied by the yolk glands in histological sections of control (*eGFP), APC* and *ß-Catenin-1(RNAi)* animals. Top: Bar graph representation for two individuals per treatment (dots). Error bars denote the SD of the mean. Bottom: Representative views of Mallory-stained sagittal sections. Yolk glands are outlined in white. Zooms (red frame) show detail. **h**, Identification of the major yolk protein species in *Smed*. Coomassie-stained SDS-PAGE of equal amounts of protein extracts from yolk capsules and individuals of the indicated *Smed* strains. White box: Major yolk protein. **i**, Fluorescent Western blot of the indicated protein extracts, probed with the custom-produced monoclonal antibody clone EO95. Loading control: Histone H3. **j**, Specificity analysis of EO95 in protein extracts of *Smed* sexual strain individuals treated with RNAi against the indicated *ferritin* genes. Asterisk: Specific EO95 signal loss under *ferritin-C(RNAi)*. **k**, Quantification of FERRITIN-C and *ß*-CATENIN-1 amounts in lysates from control (*eGFP), APC* and *ß-Catenin-1(RNAi)* sexual strain individuals via EO95 or G78 immuno-blotting. Error bars denote the SD of the mean of four technical replicates (dots) within one biological replicate; samples were normalised to the signal levels in control. Dots above error bars are data points and do not represent statistical significance. **l**, Colorimetric whole-mount in situ hybridisation expression patterns of the four yolk ferritins in sexual *Smed* under control (*eGFP), APC* and *ß-Catenin-1(RNAi)*. Scale bar: 1 mm unless otherwise noted.

This view also entails that a positive pleiotropy of Wnt signalling in egg-laying reproduction might explain the repeated selection of Wnt-dependent regeneration defects. So far, the functions of planarian Wnt signalling have been mostly studied in the asexual strains of the model species *Smed* and *D. japonica*, which never develop the various components of the planarian hermaphroditic reproductive system (RS; e.g., testes, ovaries, yolk glands, and the copulatory apparatus) under standard laboratory conditions. To explore putative Wnt signalling functions in the planarian RS, we turned to the sexual laboratory strain of the model species *Smed*. The RS in adults of this strain^64,65^ (Fig. 5c) can be collectively ablated through the knockdown of the orphan G-protein coupled receptor *ophis*^66^. As a proxy for a putative association between Wnt pathway activity and the RS, we quantitatively compared ß-CATENIN-1 levels between the sexual and asexual *Smed* strains using our previously established quantitative Western blotting methods^17^ (Fig. 5d). Interestingly, Wnt signalling levels were consistently higher in the sexual strain as compared to the asexual strain, and *ophis(RNAi)* reduced ß-CATENIN-1 levels to asexual-like levels, thus indicating significant canonical Wnt signalling activity in association with the RS. Consistently, we detected the expression of several Wnt receptors and other pathway components in multiple RS-associated tissues (Fig. 5e, S. Fig. 5b,c). To probe the functional requirements of Wnt signalling in the RS, we globally activated or inhibited Wnt signalling in mature egg-laying animals via *APC(RNAi)* or *ß-Catenin-1(RNAi)* and observed the ensuing consequences via our panel of RS component marker genes (Fig. 5f, S. Fig. 5d). Of note, the experimental animals were examined before morphological indications of anteriorization or posteriorization that are elicited by prolonged RNAi treatments^17^. The morphology of testes and ovaries appeared overtly unaffected under our experimental conditions, although some abnormalities in testes lobe morphology were apparent in *APC(RNAi)* (S. Fig. 5d). In sharp contrast, the expression of both yolk and shell gland markers was practically undetectable under *ß-Catenin-1(RNAi)* and the yolk gland marker further often appeared denser in *APC(RNAi)* (Fig. 5f, S. Fig. 5e). Consistently, the quantification of the relative crossectional area of the yolk glands in histological sections of mature control, *ß-Catenin-1* and *APC(RNAi)* animals revealed the near-complete ablation of the yolk gland by *ß-Catenin-1(RNAi)* (Fig. 5g), thus demonstrating the overt Wnt-dependence of yolk gland maintenance in *Smed*.

Given the critical importance of yolk as a life history parameter, we sought to develop a quantitative assay for directly measuring yolk content. An acrylamide gel analysis of yolk revealed a prominent 25 kDa protein species, which was also detectable in sexual strain extracts, but not in asexual strain extracts (Fig. 5h, S. Fig. 5f). Mass spectrometry identified the major constituents of the bands as four *Smed FERRITINS* (here named A to D), which were specifically expressed in the sexual *Smed* strain in expression patterns highly reminiscent of the generic yolk gland marker *surfactant-b*^67^ (S. Fig. 5g). FERRITINS are therefore the major yolk protein in *Smed*, which is consistent with recent findings in the planarian species *Dugesia ryukyuensis*^68^. Next, we raised a monoclonal antibody against the FERRITIN pool eluted from gel slices. The clone EO95 specifically recognised the ~ 25 kDa FERRITIN band (Fig. 5i), and lysates of individual *ferritin RNAi* treatments established its likely specificity for FERRITIN-C (Fig. 5j). To quantify the dependence of yolk production on Wnt signalling levels, we compare FERRITIN-C amounts in adult sexual *Smed* under reduced (*ß-Catenin-1(RNAi)*) or increased (*APC(RNAi)*) Wnt pathway activity via quantitative Western Blotting. *ß-Catenin-1(RNAi)* resulted in a near-complete loss of FERRITIN-C (Fig. 5k). The whole mount expression patterns of the four yolk ferritins were also strongly depleted by *ß-Catenin-1(RNAi)* and possibly slightly increased in *APC(RNAi)* (Fig. 5l). Overall, our results demonstrate so far unknown roles of Wnt signalling in the RS, including a functional requirement in the maintenance of shell and yolk glands and quantitative influence on yolk content in the model species *Smed*.

Our results are, therefore, consistent with the following working model (Fig. 6a, centre): Robust whole-body regeneration in planarians is necessary for fissiparous reproduction but dispensable for reproduction via egg-laying (centre). Each reproductive strategy has different costs and benefits, and natural selection favours individuals with traits best adapted to the specific environment (Fig. 6a red arrows). Fissiparous (asexual) reproduction avoids the cost of males and allows the establishment of a new population from a single individual ^69,70^, which may be particularly advantageous for the rapid colonisation of temporary habitats or fast-flowing mountain streams. Egg laying and the commonly entailed meiotic recombination generate new allele combinations that may benefit long-term population survival in stable habitats, e.g., large lakes or the sea. Selection for each of the two reproductive strategies entails opposite selective pressures on planarian Wnt pathway activity (top): While reproductive performance via egg-laying is positively influenced by high Wnt pathway activity (e.g., yolk production in *Smed*), excess pathway activity interferes with whole-body regeneration and, thus fissiparous reproduction. Hence the model envisages the emergence of planarian regeneration defects via the habitat-specific selection for increased yolk content or larger egg capsules and concomitant elevation of Wnt signalling levels to a point where they begin to interfere with head regeneration. The many egg-laying and whole-body regeneration competent species (e.g., the sexual *Smed* strain) are envisaged to occupy intermediate levels of the trade-off regime, i.e. intermediate Wnt signalling levels and intermediate investment in sexual reproduction that do not yet interfere with regeneration.

**Figure 6.**
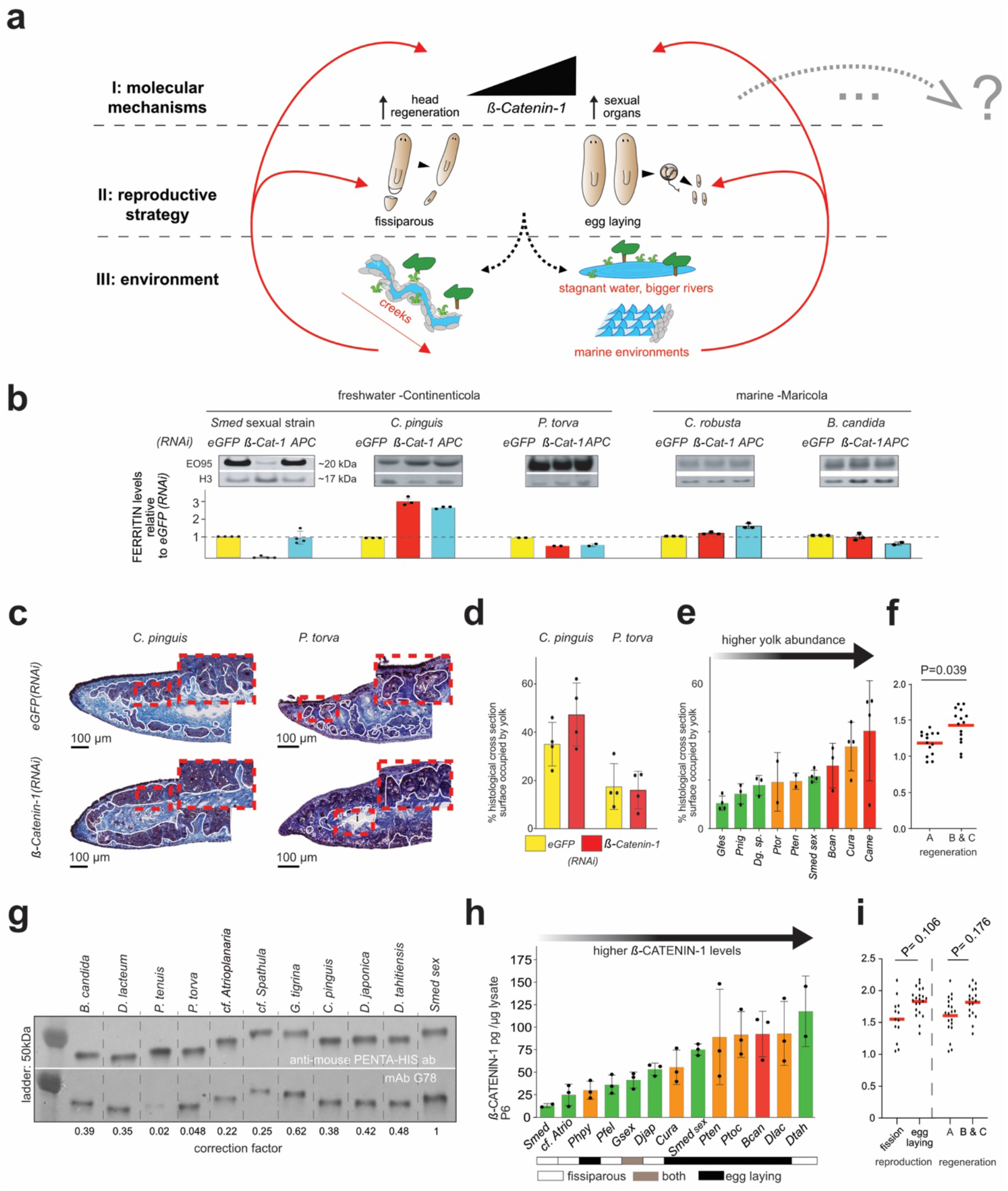
Evolution of Wnt-dependent regeneration defects in planaria. **a**, Multi-scale model motivated by our data. See text for details. **b**, Test of Wnt-dependence of yolk production in other planarian species. Top: Quantitative Western blot analysis of FERRITIN-C (EO95 Ab) and H3 (loading control) in lysates of the indicated species and RNAi conditions. Bottom: Band intensity quantification relative to control (*eGFP*). Error bars denote the SD of two to four technical replicates (dots) within one biological replicate (bottom). **c**, Yolk gland cross-sectional area analysis in sagittal sections of control *eGFP* and *ß-Catenin-1(RNAi)* treated *Cura pinguis* and *Planaria torva*. Yolk glands are outlined in white, zooms (red frame) show details. **d**, Quantitative analysis of the data in c. Error bars denote the SD of the mean; dots analysed individuals. **e**, Correlation between relative yolk gland cross-sectional area and regenerative abilities in representative egg-laying species. Quantifications in the indicated species as in c, d. Species are ordered by relative yolk content and color-coded according to head regeneration abilities (Group A/robust regeneration: green; Group B/restricted regeneration: orange; Group C/poor regeneration: red). Error bars: SD of the mean. Dots denote analysed individuals. **f**, Statistical analysis of the data in e, re-plotted as log10 of the mean yolk content of individual measurements in Group A versus Groups B, C. Statistical significance of the difference was assessed via Linear Mixed Model (LMM)_-and the obtained P-value (0.0039) indicates a significant difference between the two distributions. **g**, Calibration of the anti-*ß*-Catenin-1 mAB G78 signal via quantitative Western blot analysis of recombinant His-tagged *ß*-CATENIN-1 fragments covering the conserved armadillo repeat region of *Smed-ß-*CATENIN-1 (antigen against which G78 was raised). The ratio between the quantified penta-His and G78 signal of each fragment (bottom) was used as a correction factor for species-specific differences in G78 affinity. Molecular weight markers and species as indicated. **h**, Correlation between Wnt pathway activity, regenerative abilities and reproduction mode across the species collection. *ß*-CATENIN-1 levels in the tail tip (piece 6) were quantified as a proxy for canonical Wnt signalling activity via quantitative Western blotting with the G78 mAB as in Fig. 4a, b. Ordering of the indicated species or strains by measured *ß*-CATENIN-1 levels and colorcoding according to head regeneration abilities as in e. The predominant reproduction mode is indicated below. Error bars denote the SD of the mean for two to three biological replicates, each with two to four technical replicates. Dots denote individual biological replicates. **i**, Statistical analysis of the data in f, re-plotted as log10 of the mean *ß*-CATENIN-1 tail tip levels in fissiparous vs egg-laying strains/species (left) or robust regeneration vs restricted or poor regeneration (Groups A vs B, C). Statistical significance of the difference was assessed via Linear Mixed Model (LMM), and the obtained P-values indicate a non-significant difference between the two distributions in both analyses.

The model makes several predictions: First, the Wnt dependence of yolk production should be conserved across planarian phylogeny. Second, amongst egg-laying species, regeneration-restricted species should generally invest more in reproduction than robust regenerators. Third, Wnt signalling levels across planarian phylogeny should positively correlate with egg-laying versus fissiparous reproduction and peak in regeneration-deficient species. Our species collection and the tools established during this study allowed a first experimental exploration of these predictions. To ask whether the Wnt dependence of yolk production is conserved across planarian phylogeny, we first screened our FERRITIN-C monoclonal antibody panel for crossreactivity with cocoon lysates of different planarian species and found that clone EO95 identified a band of the expected size in egg-laying adults of *C. pinguis, P. torva, C. robusta and B. candida* (but not in *P. tenuis;* data not shown). Moreover, the band was barely detectable in lysates of sexually immature *C. pinguis* or *P. torva* hatchlings (S. Fig. 6b), which further confirms the utility of the antibody signal as a proxy for the yolk content of different planarian species. In sharp contrast to the previously established Wnt signalling dependence of FERRITIN content of sexually mature *Smed*, we found that the previously validated *APC* or *ß-Catenin-1(RNAi)* treatments of our species panel failed to elicit striking changes of the EO95 signal (Fig. 6b). To exclude the possibility of lineage-specific changes in the ferritin gene complement of the yolk, we additionally quantified the yolk gland crossectional area in histological sections of *ß-Catenin-1(RNAi)* treated *C. pinguis and P. torva*. As shown in Fig. 6c-d, both species maintained their yolk glands under *ß-Catenin-1(RNAi)*, despite significant reductions of *ß-CATENIN-1* levels under the experimental conditions (Fig. 4b). While the Wnt signalling dependence of yolk production is therefore unlikely to be deeply conserved across planarian phylogeny, the possibility remains that other positive pleiotropies of Wnt signalling in the RS (e.g., shell gland specification or testes-specific functions) mediate the predicted positive association between Wnt pathway activity and the RS in specific lineages.

To explore the predicted correlation of relative investment in sexual reproduction with regeneration defects, we quantified the relative cross-sectional area of yolk glands as a proxy. In the nine collection species that consistently produce egg capsules under our laboratory conditions, the yolk glands of restricted or poor regeneration species (groups B/C) indeed tended to occupy a larger fraction of the cross-sectional area as in regeneration competent species (group A; Fig. 6e). This tendency was statistically significant despite substantial inter-animal variations in the yolk content in some species (e.g., *P. tenuis, P. torva* or *Camerata robusta*) (Fig. 6f), which may reflect non-synchronous reproduction cycles under our laboratory culture conditions. These data are further consistent with published reports of extraordinary investments in reproduction in the restricted regeneration species *D. lacteum* or *P. torva*^71^, including the continuation of egg production even under starvation and extraordinarily large egg capsules in the case of *D. lacteum*. Overall, these data are consistent with the postulated higher investments in egg-laying reproduction in regeneration-deficient species compared to regeneration-competent ones.

Finally, to probe the predicted correlation between Wnt signalling levels and reproductive strategy or regeneration deficiencies, we calibrated our G78 pan-planarian anti-*ß*-CATENIN antibody as a proxy for inter-species comparisons of Wnt pathway activity. Using known amounts of *in vitro* translated recombinant armadillo repeat fragments of the different species as quantitative Western blotting standards, we were able to measure and compare absolute ß-CATENIN-1 amounts in the lysates of different species (Fig. 6g). Our species panel for this experiment included a total of 13 collection species that represented a broad sampling of regenerative abilities, reproductive strategies and phylogenetic position (Fig. 6h). All species except *C. robusta* (S. Fig. 6a) displayed the posterior to the anterior gradation of ß-CATENIN-1 concentration, consistent with the likely conservation of Wnt-gradient mediated A/P-patterning^17^ across planarian phylogeny (not shown). ß-CATENIN-1 amounts in the tail tip samples of the different species varied by more than 10-fold (Fig. 6h), suggesting significant variations in signalling gradient amplitudes between species. Interestingly, restricted or poor regeneration species were enriched amongst the species with the highest ß-CATENIN-1 levels. The statistical analysis of tail tip *ß*-CATENIN-1 levels relative to reproductive strategy (fissiparous versus egg-laying) or regenerative abilities (robust versus restricted and poor) similarly indicated a tendency towards higher ß-CATENIN-1 levels in egg-laying and regeneration impaired species, yet without reaching statistical significance (Fig. 6i). The latter could be largely attributed to the *D. tahitiensis*, in which we measured the highest *ß*-CATENIN-1 levels of any of the species tested, even though the species displays robust whole-body regeneration and exclusively fissiparous reproduction under our laboratory conditions. Although the *ß*-CATENIN-1 quantifications in the present species panel failed to unequivocally support the association between Wnt signalling levels, regeneration and reproduction, our model nevertheless remains a useful working hypothesis for guiding the future analysis of broader species samplings or cell biological analyses of Wnt signalling mechanisms in individual species.

## Discussion

Overall, our study cannot authoritatively answer the question of why some planarians regenerate while others cannot. In fact, such a single or simple answer may not exist in the face of the deep splits between planarian lineages and the associated divergence of molecular mechanisms that we discovered (e.g., RNAi-susceptibility or the *ß-Catenin-1*-dependence of yolk production). Our model of functional pleiotropies between Wnt signalling, regeneration and reproduction and selection of regeneration as a correlate of fissiparous reproduction provides a useful working hypothesis for the further pursuit of a holistic understanding of planarian regeneration. Observations consistent with the hypothesis include the likely frequent transitions between robust and restricted regeneration in planarian phylogeny (Fig. 3), the experimental “rescue” of independently evolved head regeneration defects via the experimental inhibition of Wnt signalling (Fig. 4), multiple functional requirements of Wnt signalling in the reproductive system of the model species (Fig. 5) and tentative indications of positive correlations between Wnt signalling levels, regeneration and reproduction across the widely diverged planarian species (Fig. 6). On the mechanistic level, the assumed pivotal importance of Wnt signalling is plausible on the basis of the established pathway functions as both a molecular switch during head/tail regeneration and as provider of positional identity to the adult pluripotent stem cells in non-regenerating intact animals (neoblasts)^17–19^. Neoblasts generate and re-generate all planarian cell types^14^, including in the reproductive system^72^. Hence an elevation of the somatic Wnt signalling gradient amplitude might simultaneously upregulate yolk or testes formation, while also challenging the downregulation of pathway activity soon after wounding as a necessary prerequisite of head regeneration. More generally, our pleiotropy model predicts that regeneration and reproduction are under the influence of a common Wnt signalling source. Although our RNAi experiments are consistent with this premise, the lack of organ-selectivity of the protocol leaves open the possibility of organ-autonomous Wnt signalling in the reproductive system and, therefore, functional uncoupling between reproductive pathway functions and regeneration. Similarly, the model assumes a functional excess of steady-state Wnt signalling as a common cause of regeneration defects. Though again plausible on the basis of the *ß-Catenin-1(RNAi)*-induced rescue of head regeneration defects across planarian phylogeny, an alternative interpretation of these data is that Wnt inhibition constitutes a deep developmental constraint in the neoblast-mediated formation of the planarian head. Accordingly, Wnt inhibition might be able to override all upstream physiological control mechanisms, inclusive of potentially Wnt-independent causes of head regeneration defects, and thus bypass, rather than rescue head regeneration defects.

Above all, our working model stresses the need for a more fine-grained understanding of the cellular sources and targets of Wnt signalling, both during head regeneration and in the formation and maintenance of the reproductive system. In addition, a similarly fine-grained understanding of the mechanistic causes of head regeneration failures in regeneration-deficient species will be necessary. The deep splits between planarian lineages and the lack of conservation of Wnt-dependence of yolk production beyond *S. mediterranea* indicate that results obtained in one planarian species may not necessarily apply elsewhere in the taxon, with the rapid diversification of molecular mechanisms in *Caenorhabditis* species as a case in point^73^. A further core premise of our working model is that Wnt signalling activity is the target of natural selection. Clearly, this premise cannot be tested on laboratory populations and will require dedicated field studies in natural populations. While illustrating the general challenge of linking proximate to ultimate mechanisms in evolution, these considerations highlight the experimental opportunities that planarians and specifically our species collection offer to regeneration research.

## Acknowledgements

We thank Dr Helena Bilandzija, Dr Mette Handberg-Thorsager Dr Salvator D’Aniello, Dr Tom Stückemann and Dr Tobias Boothe for fieldwork support. We thank Heino Andreas, Rick Kluivert, Til Schubert and our student helpers for worm care and species collection maintenance. We thank Dr Jun-Hoe Lee and Dr Ian Henry for support with phylogenetic analyses, and Dr Tobias Boothe for imaging support. We thank the following facilities for their support: the Center for Molecular and Cellular Bioengineering (CMCB) Histology facility, the DRESDEN-concept Genome Center and the CMCB Light microscopy facility. We thank the following MPI-CBG facilities for their support: the Protein Expression and Purification Facility, the Antibody Facility, and the Mass Spectrometry Facility. We thank Rink lab members for lively discussions. This project received funding from the European Research Council (ERC) under the European Union’s Horizon 2020 research and innovation program (grant agreement number 649024), from the German Research Foundation (project RI 2449/51), and from the Max Planck Society funding (J.C.R.). JEJR acknowledges funding from the NHMRC Investigator Grant (#1177305) and the Sir Zelman Cowen Foundation.

## Author Contributions

M.V. and J.C.R. conceptualised the study. M.V., J.C.R, J.C., C.P., B.E., M.G., J.E.J.R., H. T-K. V. and F.C. collected samples. M.V., M.I., J.C., S.K., M.G. and A.G. performed the experiments. A.R., J.N.B., F.T., and C.B. performed the data analysis for the phylogeny. M.V. and J.C.R. wrote the manuscript. All the authors contributed to the revision of the manuscript text.

## Methods

### Field collections

The MPI-NATplanarian collection was assembled via dedicated field expeditions, mostly prior to 2015. Planarians were collected from the underside of stones, aquatic plants or other submerged objects with the help of a brush. Trapping was also used, employing submerged plastic containers with an appropriately perforated lid and baited with liver (S. Fig.1a). The collected planarians were transferred to 50 ml Falcon tubes, maintaining low animal densities to avoid worm lysis. Tubes were kept cold, and daily water changes with water from the collection site were carried out throughout the duration of the field campaign. The GPS coordinates and habitat features of collection sites were recorded in a dedicated database.

Prolecithophoran specimens were sampled from brown algae in the Adriatic sea in a harbour on the island of Krk, Croatia. The algal samples were first incubated for at least 15 min in a 1:1 7.14% MgCl_2_ × 6H_2_O and seawater, and the solution was then filtered through a 63 μm pore-sized mesh. All animals retained in the mesh were washed with seawater into a Petri dish, and prolecithophorans were collected with Pasteur pipettes under a stereo microscope. Until use for experiments, prolecithophorans were maintained according to Grosbusch *et al*. (2019).

Species can be made available to members of the community upon request to the corresponding author, subject to availability.

### Animal husbandry

To reduce the chance of inadvertent pathogen introductions, field-collected planarians were initially treated for 1-3 days with a cocktail containing the following reagents diluted in the appropriate species-specific culture media (final concentrations): rifampicin at 100 ug/ml (stock diluted in DMSO), erythromycin 4 ug/ml, gentamycin 50 ug/ml, vancomycin 10 ug/ml, ciprofloxacin 4 ug/ml, colistin 20 ug/ml, Penicillin/Streptomycin/Amphotericin B (Millipore 516104) 1:100 from stock, and Spirohexol Plus 250 (JBL;1007100) (12 μl from stock per 10 ml of final solution). To establish long-term cultures of field-collected animals, one of three water formulations (see below) and three culture temperatures (10, 14, or 20 °C) were chosen as starting point according to the field notes. “Montjuïc planarian water”, the widely used salt solution for *Smed* laboratory cultures^74^, was used for most planarian species. “*Polycelis felina* water” (PofW) is the 1X combination of two commercial aquaria salts, DRAK Aquaristik GH+ (DRAK-Aquaristik GmbH, Germany, W0060100) and KH+ (DRAK-Aquaristik GmbH, Germany, W0070100). 20x stock solutions of both products were mixed and diluted to the final 1x concentration of each salt with distilled water. Afterwards, 0.25 ml/L Tetra ToruMin (Tetra GmbH, Germany) was added to mimic organic solutes. Marine water was prepared by diluting Classic Meersalz (Tropic Marin, Germany, 10134) in MilliQ water to a final salt concentration of 32 g/l with the help of a refractometer. For species cultured at 10 or 14 °C, the water formulations were always pre-cooled to minimize temperature shocks. To establish suitable food sources for wild-collected planarian species, small-scale feeding trials with organic calf liver paste, frozen and irradiated (60GY) mealworms (*Tenebrio molitor*) or rinsed *Drosophila* larvae were carried out and the food source eliciting the strongest feeding response was chosen as sustenance food. For *Smed*, the culture media was supplemented with gentamycin sulfate at 50 ug/ml. Episodic gentamycin supplementation was also used for other species in case of indications of poor culture health. Collection species were maintained in plastic dishes and fed at intervals with the sustenance food of choice for the species. Feeding was generally coupled with cleaning/rinsing with temperature-equilibrated water formulations. Feeding/cleaning intervals were adjusted depending on culture temperatures and feeding schedules.

### Live imaging

Flatworms were imaged with a Nikon AZ100M microscope equipped with a Digital Sight DS-Fi1 camera or a ZEISS Stereo Microscope Stemi 508 equipped with a Digital ZEISS Axiocam 208 colour camera. Only animals from healthy laboratory populations were chosen for the experiments. Prolecithophorans were imaged either with a Leica DM 5000 B microscope equipped with a Leica DFC 490 camera or on a Leitz Diaplan light microscope equipped with a DFK 33UX264 camera.

### Quantitative analysis of head regeneration

To quantify species-specific head regeneration abilities, 7 to 20 specimens per strain were cut into six even pieces along the anteroposterior body axis. For species ~5 mm in length, animals were cut into four pieces (*Procedores littoralis, Procerodes plebeius* and *Camerata robusta*). Cuts were performed with the help of a microsurgical knife under a stereoscope. Animals were cold-immobilized on a wet sheet of filter paper using custom-built Pelletier cold blocks. *Dugesia sicula*, a temperature-sensitive species, was cut at room temperature. The resulting amputation fragments were grouped according to body axis position and maintained in Petri dishes or small plastic boxes in the accustomed maintenance medium of the species. Water changes were carried out daily for the first three days post-amputation and every 3-4 days for the remainder of the experiment. Eye regeneration as a morphological marker of head regeneration was scored at 3-4 day intervals for a maximum of eight weeks to account for species-specific variations in regeneration rates. Head regeneration at each position was quantified as the fraction of fragments (either initial number or surviving pieces) that successfully regenerated eyes. For Prolecithophora, the experimental setup for mid-body amputations and monitoring of regenerates is described in detail in Grosbusch *et al*. 2022^42^.

### Transcriptome sequencing and assembly

RNA extraction and quality control were performed following an established protocol^17^ and processed for 100 or 150-bp paired-end Illumina sequencing. Double-indexing was used to minimise the cross-contamination of transcriptomes. Transcriptome assembly was carried out with our established pipeline^46^. Assembly completeness was assessed using BUSCO^47^ (Benchmarking Universal Single-Copy Orthologs - v 5.2.2 - metazoa odb10 - parameters: - protein). Transdecoder (version: 5.5.0, parameters: --single_best_orf) was used for the in silico translation of the transcriptomes as input for BUSCO.

### Orthology inference and phylogenetic tree inference

An initial set of homologous groups were identified by applying OrthoFinder^51^ (version 2.5.4, parameters: -M msa -I 1.5) to all proteomes (i.e., *in silico*-translated transcriptome assemblies). The resulting homologous groups were subsequently aligned using MAFFT^75^ (version 7.487, parameters: --localpair --maxiterate 1000), and for each alignment, a gene tree was constructed using FastTree^76^ (version 2.1.10). The alignments and their corresponding phylogenetic trees were provided as input to the tree-based orthology inference program PhyloPyPruner (Thalén et al. in prep., version 1.2.3, parameters: --min-len 150 –trim-lb 4 – min-support 80 –prune MI –min-taxa 65 –min-otu-occupancy 0.0 –min-gene-occupancy 0.0) (github.com/fethalen/phylopypruner). The optimal parameters for PhyloPyPruner were chosen by comparing the outcome when adjusting for minimum sequence length, long branch trimming factor, minimum support value, minimum number of taxa, minimum OTU occupancy, tree pruning method, and minimum gene occupancy. The optimisation script, including the tested parameter values, can be found in the supplementary material. Phylogenetic trees were constructed using IQ-TREE^77^ (version: 2.1.2, parameters: -m MFP -bb 1000 -bnni) or via ASTRAL^78^ (version 5.7.1), using standard parameter settings (S. Fig. 3a). The phylogeny combining triclads, mammals and nematodes was built following the same approach as for the planarian phylogeny.

### Ancestral state reconstruction (ASR)

ASR of regeneration ability was performed on the maximum-likelihood phylogeny. First, we calculated an ultrametric phylogeny with root depth 1 using the penalised likelihood method^79^ implemented in the software TreePL^80^. Then the appropriate transition matrix for ASR was determined by fitting MK-models with equal transition rates (ER), with symmetric transition rates (SYM), and with all transition rates different (ARD) and then evaluating the model fit using the corrected Akaike information criterion (AIC). SYM was the preferred model. Next, we used stochastic character mapping^81^ implemented in the R package phytools^82^ to simulate 10.000 character histories of regeneration using the transition matrix from the best-fitting MK-model. The maximum likelihood implementation of the method was used, which samples histories from the most likely transition matrix. 10.000 iterations of burn-in, followed by 100.000 iterations where we retained every 10th character history, were performed. Finally, the transitions were summarised as the arithmetic mean of the number of changes.

To control for the species with population-dependent variation in head regeneration ability, *Girardia dorotocephala* and *Polycelis sapporo*, we created two species datasets. Those two species are categorised in group A in dataset A and group B in dataset B (see S. Fig. 3c). Figure 3e shows the analysis for dataset A.

### Identification of *ß-Catenin-1* and *APC* in multiple planarian species

Homologues of *Smed ß-Catenin-1* and *APC* in other species were identified using reciprocal BLAST against the respective transcriptome assemblies. For the assembly of the *ß-Catenin-1* phylogeny, transcript sequences were translated using transdecoder (v 5.5.0), (https://github.com/TransDecoder), and the single best ORF per sequence was selected. Translated sequences were aligned using MAFFT (v 7.490)^75^, using “--maxiterate 1000 --localpair” as parameters. Next, trimAl (v 1.4.1)^83^ was used with “-automated1” and the tree was built using IQ-TREE (v 1.6.12)^77^ with the parameters “-m MFP -bb 1000 -bnni”.

### Cloning and *RNA*-mediated gene silencing

For gene cloning, cDNA was synthesised using the SuperScript^®^ III Reverse Transcriptase (LifeTechnologies, 18080093) according to the manufacturer’s recommendations, followed by an *E. coli* RNase H step. DNA templates were amplified from cDNA using either published primers or primers designed using our transcriptomes (S. Table 4) and cloned into the pPR-T4P vector by ligation-independent cloning^84^ or the paff8cT4P vector (for recombinant protein production)^85^. For *Smed, RNAi* of specific target genes was carried out by feeding liver paste mixed with in vitro synthesised dsRNA^86^. For *RNAi* experiments in species other than *Smed, Artemia* paste obtained via sonication of *Artemia* larvae was added to the liver paste (10% of the final *RNAi* food volume) to improve feeding efficiency. *RNAi* feedings were performed every third day, with a final dsRNA concentration of 1 or 2 μg/μl food. Animals received three feedings unless stated otherwise.

### Gene expression analyses

Riboprobe production, animal fixation, colorimetric in situ hybridisation, FISH and mounting were performed largely as described^87^, but incorporating several optimisations for large planarians (> 1cm; Vila-Farré et al., in press). Representative specimens from colorimetric whole-mount in situ hybridisation were imaged with a Nikon AZ100M microscope equipped with a Digital Sight DS-Fi1 camera. For the documentation of FISH, we used a Zeiss Axio Observer.Z1 with a confocal Zeiss LSM 700 scan head equipped with a 20 or 25x objective. Brightness/contrast and color balance were adjusted using Adobe Photoshop Creative Cloud 2018 and always applied to the whole image. Figure panels were assembled using Adobe Illustrator Creative Cloud 2018 and Affinity Designer. For planarian live image montages (e.g., Fig. 1), the animals were cropped out of the original image frame and pasted onto a uniform black background.

### Identification of *Smed* yolk proteins

Two-day old egg-capsules were homogenised in 200 μl cold RIPA buffer (150 mM NaCl, 50 mM Tris-HCl pH 8.1, 0,1% SDS (w/v), 1% Sodium Deoxycholate, 1% NP-40) with a pestle and pellet mixer and incubated for 20 min on ice. Laemmli buffer (6x Laemmli buffer; 12% SDS; 0,06% Bromophenol blue; 50% Glycerol; 600mM DTT; 60mM Tris-HCl pH6.8) was added to a final concentration of 1x and incubated for 10 min at 95°C. 6 volumes of 1x Laemmli buffer were added, and 10 μl of the sample was used for SDS-PAGE (S. Fig. 5f). The bands in the gel were cut, homogenised, and analysed via mass spectrometry by the MPI-CBG mass spectrometry facility.

### In vitro transcription-translation and antibody production

The Expressway™ Maxi Cell-Free *E. coli* Expression System was used according to the manufacturer’s recommendations for the in vitro production of species-specific ß-CATENIN-1 standards. As a template, we used the ß-*Catenin*-1 fragments of different species cloned into the paff8cT4P vector. The concentration of the resulting His-tagged ß-CATENIN-1 fragments was quantified via quantitative Western blotting with a Penta-His antibody (Qiagen) and a recombinant His-tagged protein as external standard.

The anti-FERRITIN antibody EO95 was raised against a pool of endogenous proteins eluted from gel slices following established protocols and procedures of the MPI-CBG Antibody Facility^17^.

### Quantitative Western blotting (qWB) and qWB analysis

qWB was performed as described in^17^ with minor modifications. Animals were fixed in zinc fixative (100 mM Zn Cl2 in 100% EtOH) for 30 min at 4°C and then stored at −80°C. To prepare the complete lysis buffer, to 1 ml of freshly thawed 9 M Urea lysis buffer (9M Urea, 100mM NaH2PO4, 10 mM Tris, 2% SDS, 130 mM DTT, 1 mM MgCl2, pH 8.0) we added phosphatase inhibitor (25 μl of 40X PhosSTOP), benzonase (10 μl of 250 Unit/ml solution, final conc.: 1%) and protease inhibitor cocktail (20 μl of 100X HaltTM Protease Inhibitor Cocktail). Samples were run out in technical quadruplicates on NuPAGE Novex 4–12% Bis-Tris protein gels in 1x MES-SDS running buffer, transferred onto nitrocellulose membranes in 1X NuPage Transfer Buffer (25 mM Bicine, 25 mM Bis-Tris (free base), 1 mM EDTA, 0.05 mM chlorobutanol, 1mM NaHSO3, 0.01% SDS, 20% methanol, pH 7.2), blocked in 5% soya protein powder solutions in 1x PBS and incubated with primary antibody (anti-*Smed*-ß-CATENIN-1 antibody G78 mouse monoclonal at 1 μg/ml; anti-*Smed*-FERRITIN antibody EO95 mouse monoclonal at 0.1 μg/ml; anti-rabbit Histone H3 antibody (Abcam, ab1791) at 10 ng/mL) in 5% soya protein powder in 1x PBS with 0.1% Tween20. Membranes were washed with washing buffer (1x PBS with 0.1% Tween20) before incubation with infrared fluorescent secondary antibodies (anti-Mouse 770CW, LICOR and anti-Rabbit IRDye 680LT, LICOR) diluted at 1:20000 in blocking solution. Membranes were washed with washing buffer, followed by a final wash step in 1x PBS without Tween20. Stained membranes were dried and imaged on an LI-COR Odyssey imager.

All western blot image quantifications were conducted using ImageStudioLite software (LICOR). Rectangles were drawn around the protein of interest (ß-CATENIN-1 or FERRITIN) and H3 (loading control) bands, and then the background was subtracted from the total signal using the ‘‘Median” method. Each ß-CATENIN-1 signal was normalised to the corresponding H3 signal to account for technical variability.

To obtain absolute ß-CATENIN-1 concentrations (pg/ug total protein lysate) from protein samples, 200, 800, and 1500 pg of the respective recombinant ß-CATENIN-1 standard were run out on the same gels as the experimental samples. A three-point regression analysis on the quantified standard bands was, in turn, used to infer the amount of ß-CATENIN-1 in the experimental samples on the same blots. The sample concentration was calculated by normalising the measured ß-CATENIN-1 amount to the loaded sample volume (protein lysate). Unless noted otherwise, each experiment was carried out in three biological replicates (independent lysate preparations), each comprising four technical replicates (independent gels/blots of the same lysate sample).

### Histological analysis of yolk glands

Specimens were first relaxed with a 1:3000 dilution of linalool (Sigma, L2602) and subsequently killed and fixed with Bouin’s solution^88^ for 12 hours, transferred to 70% ethanol and cleared in xylene before paraffin embedding. Sectioning and staining were conducted by the Dresden Concept Histology facility. 5 μm transverse sections were attached to glass slides, stained in Mallory-Casson and mounted in DPX. All wide-field images were generated on an Axioscan Zeiss Axio Scan.Z1 slide scanner. The percentage area occupied by yolk gland tissue in images of five pre-pharyngeal cross-sections was outlined manually on the basis of the strongly contrasting yolk granules and quantified using the software Fiji^89^.

### Statistical analysis

The association between regeneration ability, reproduction mode and ß-CATENIN-1 abundance was tested using linear mixed-effects models. First, the measured ß-CATENIN-1 amounts in tail tip lysates were log10 transformed and inspected for approximately normal distribution. Then a linear mixed model was fit using the *lmer* function in the R package lme4 (v 1.1-30) in R (v 4.1.2), with regeneration ability or sexual system as the fixed effect and species identity as a random effect: *y* = *x + (1 | species)*. For significance testing, the degrees of freedom were approximated using Satterthwaite’s method as implemented in the R package *lmerTest* (v 3.1-3). The same method was used to evaluate the relationship between regeneration ability and yolk abundance.

