## Supplementary Figures and Supplementary Tables for "Probing the evolutionary dynamics of whole-body regeneration within planarian flatworms"

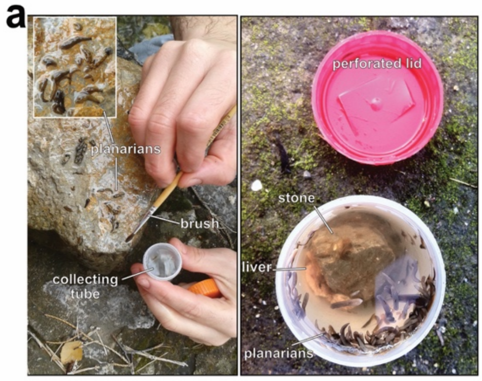


**b**

| Montjuïc planarian water (PW) | *Polycelis felina* water (PofW) | Marine water (MW) |
|---|---|---|
| 1.6 mM NaCl | 80 mg/l Duradrakon GH-Plus | 32 mg/l marine sales |
| 1.0 mM $CaCl_2*2H_2O$ | 20 mg/l Duradrakon KH-Plus | |
| 1.0 mM $MgSO_4*7H_2O$ | 0,25 ml/l Toru Min (Tetra) | |
| 0.1 mM $MgCl_2*6H_2O$ | | **17 mg/l brackish water (17 mg/l BW)** |
| 0.1 mM KCl | | dillution of MW to 17 mg/l with PW |
| 1.2 mM $NaHCO_3$ | | |
| Adjust pH to ~7.5 with 1 M HCl | | |

| Species | Water types | Temperatures °C | Food types |
|---|---|---|---|
| *Smed* | PW | 20 | liver paste |
| *Planaria torva* | PW | 10 | liver paste |
| *Polycelis felina* | PofW | 10 | liver paste/meal worm |
| *Bdelloura candida* | MW | 20 | liver paste |
| *Camerata robusta* | 17 mg/l BW | 20 | liver paste |

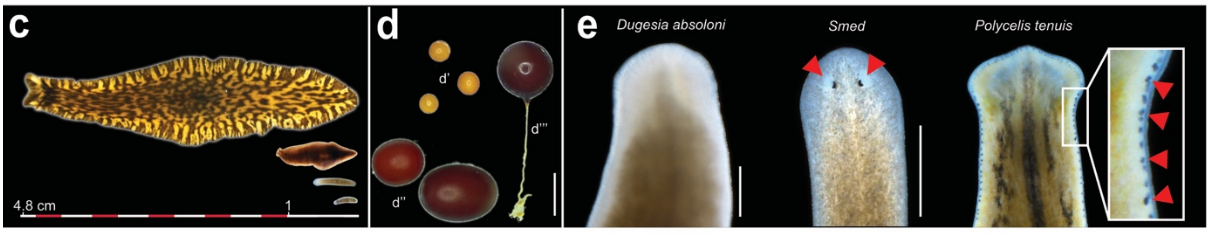


**Supplementary Figure 1. a,** Illustration of the two predominant field sampling strategies. Manual sampling with brushes (left) or simple, baited traps (e.g., liver) (right). **b,** Formulation of the three main culture media (water types) in use in the collection. The indicated components are diluted with autoclaved Milli Q water unless otherwise indicated. pH is only adjusted for “Montjuïc planarian water”. Below: representative examples of species-specific culture conditions (water type, temperature and food types). **c,** Live image montage illustrating the dramatic body size variations in different planarian species. From top to bottom: *Bdellocephala angarensis*, *Bdellocephala brunnea*, *Smed* (asexual strain) and *Camerata robusta*. **d,** Live image montage illustrating the large differences in egg capsule size and shape between different species. d’ *Camerata robusta*; d’’ *Polycelis tenuis*; d’’’ *Smed*. **e,** Live image montage illustrating the variability in eye number and placement between different species. Red triangle, eyes. Scale bar: 1 mm unless noted.

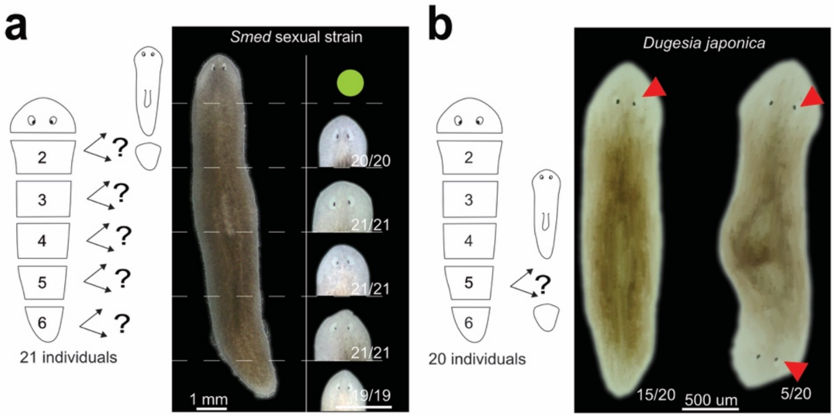


**Supplementary Figure 2. a,** Cartoon of the amputation assay and live images demonstrating robust head regeneration in sexual strain individuals of *Smed,* equal to asexual strain individuals (compare to Fig. 2a). **b,** Occasional double-head regeneration from posterior pieces in *Dugesia japonica*. The relative frequency of double head regeneration for piece 5 is indicated. Red triangles highlight eyes.

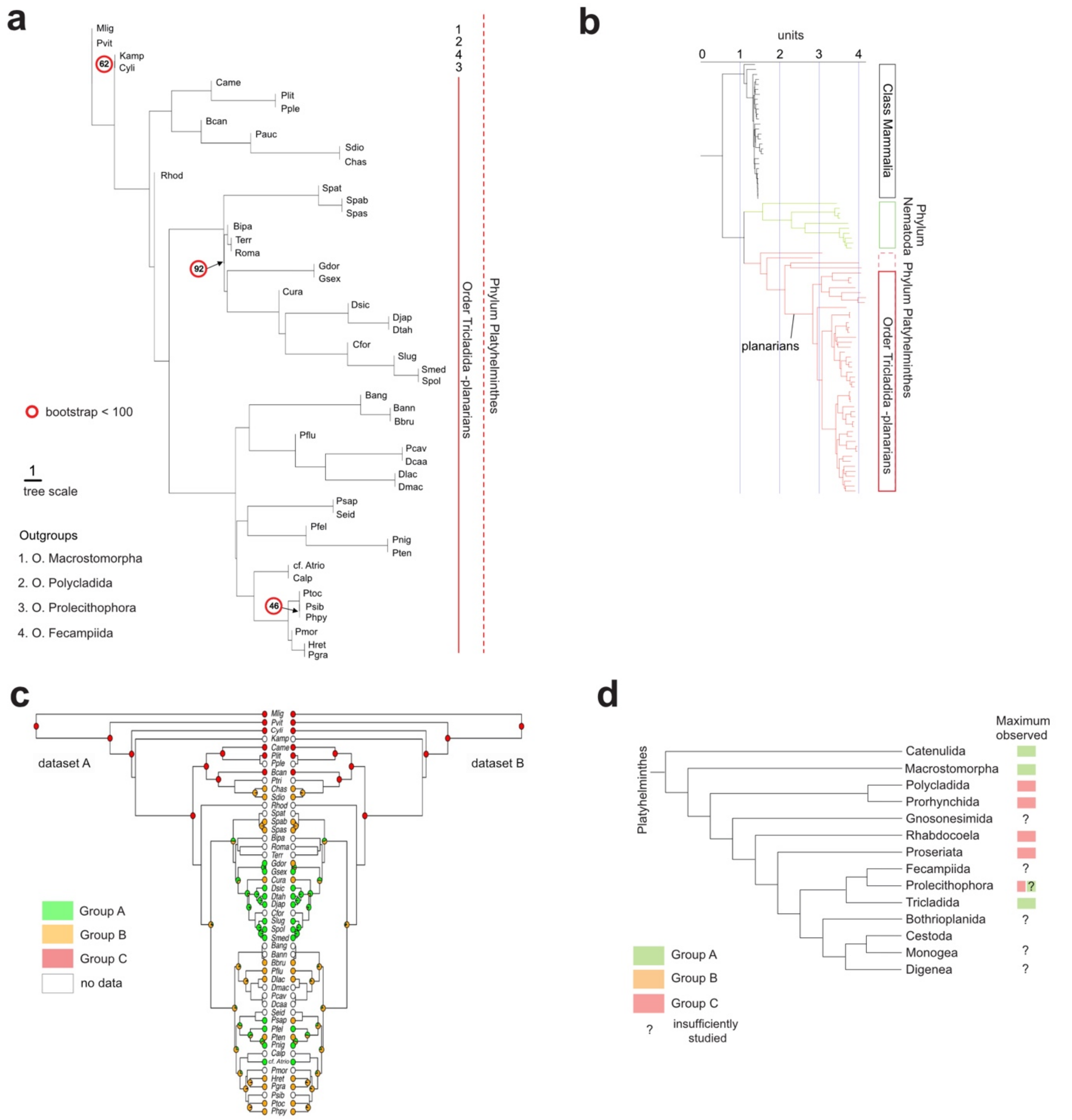


**Supplementary Figure 3. a,** Astral tree inferred from our dataset with representatives of all the suborders of the Tricladida. The tree topology is similar to that of the Maximum-Likelihood (ML) tree in Fig. 3d, thus further supporting our underlying phylogenetic hypotheses. **b**, Quantitative branch length comparison between mammals, nematodes and Platyhelminthes via Maximum-Likelihood (ML) analysis (see methods). The branch lengths of planarian clades and species (red) indicate much greater sequence divergence as compared to mammals and are at least on par with the sequence divergence amongst the nematode species included. **c**, Detailed ancestral state reconstruction inferred (see methods). To control for the species with population-dependent variation in head regeneration ability, *Girardia dorotocephala* and *Polycelis sapporo*, we created two species datasets. Those two species are categorised in group A in dataset A and in group B in dataset B. Figure 3e shows the analysis for dataset A. Data summarised in Fig. 3e corresponds to dataset A. Pie charts at nodes show the proportion of character histories with the indicated state for the capability to regenerate. **d**, Head regeneration capacity in all major free-living Platyhelminthes taxa, modified from[1,2] and adapted to our head regeneration categories.

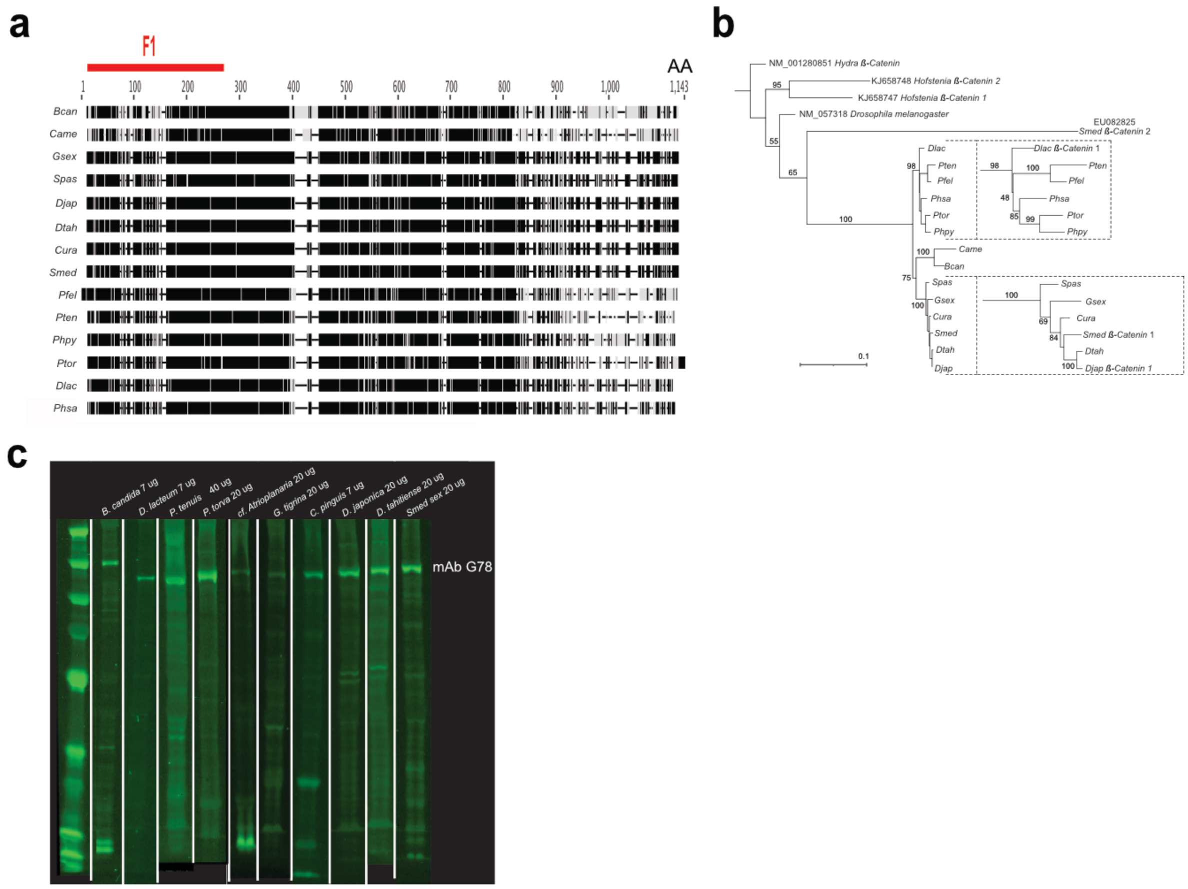


**Supplementary Figure 4. a,** Alignment of the *Smed ß*-CATENIN-1 protein sequence with the *ß*-CATENIN-1 homologues (see below) of the 13 planarian species for which the species-specific G78 affinity was calibrated. Red line: The region of *Smed ß*-CATENIN-1 against which G78 was raised. **b,** *ß*-CATENIN-1 orthologues in different planarian species. ML tree inferred from published *ß-Catenin-1* gene sequences (*Dlac*, *Smed*, *Djap*), *ß*-CATENIN-1 orthologues in the transcriptomes of the indicated species obtained by reciprocal BLAST with *Smed ß*-CATENIN-1 and *Smed- ß-Catenin-2* and several *ß-Catenin* sequences from outgroups. Dashed insets: zoom of selected branches, largely mirroring the phylogenetic distance between the species. **c,** Uncropped version of the fluorometric Western blot gel image shown in Fig. 4a, displaying the G78 anti-*ß*-CATENIN-1 antibody background reactivity in different planarian species.

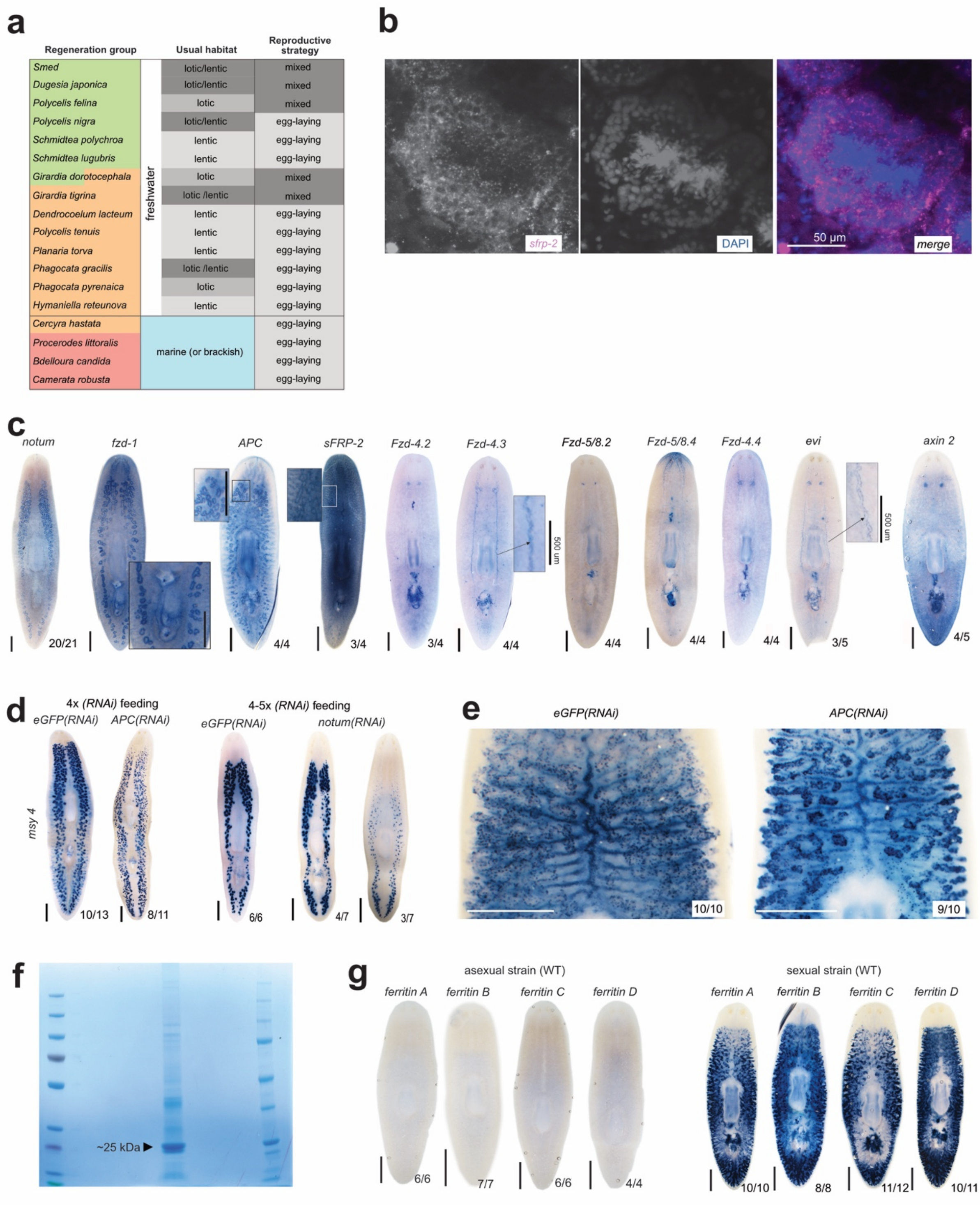


**Supplementary Figure 5. a,** Correlation between head regeneration capacity, habitat and reproductive strategy in selected species. Lentic: standing freshwater systems (e.g., ponds, lakes, vernal pools). Lotic: aquatic systems that consist of flowing fresh water (e.g., creeks, rivers, springs). "mixed" indicates that reproduction both by egg-laying and fission is known for the species. **b,** Fluorescent situ hybridisation expression patterns of *sfrp-2* in *Smed* testes. **c,** Whole-mount colorimetric (NBT/BCIP) in situ hybridisation expression patterns of selected Wnt signalling pathway genes in the reproductive system of *Smed*. Gene names as indicated. **d,** Whole-mount in situ hybridisation expression patterns of the testes marker *msy-4* after *eGFP, APC(RNAi) (left)* and *eGFP*, *notum(RNAi) (right)*. **e,** Detail of Fig. 4f, illustrating denser staining of the yolk marker *surfactant-b* whole-mount colorimetric in situ hybridisation expression pattern in *APC(RNAi)* as compared to control *(eGFP(RNAi))*. **f,** SDS-page gel of protein extracted from sexual individuals, showing the two bands analysed via mass spectrometry. **g,** Whole-mount colorimetric in situ hybridisation expression patterns of the indicated four yolk *ferritin*s in asexual (left) and sexual (right) S*med.* Scale bar: 1 mm unless otherwise noted; number pairs indicate the relative frequency of the shown pattern.

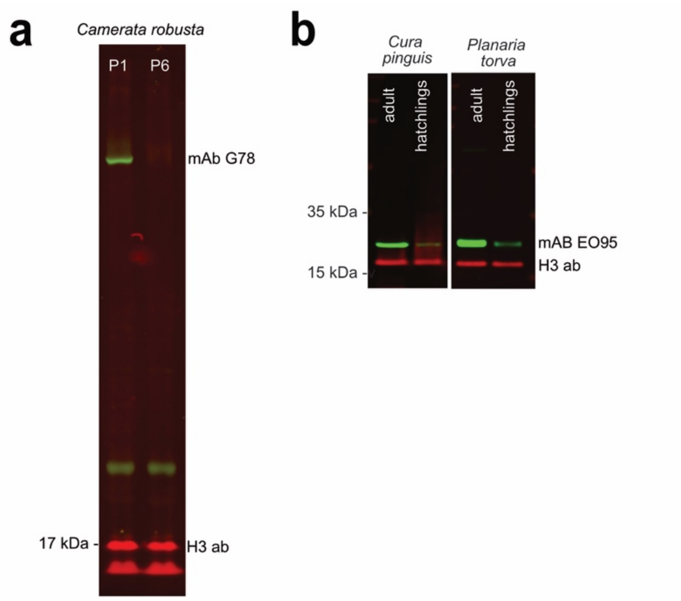


**Supplementary Figure 6. a,** Fluorometric Western blot probed with the anti-*ß*-CATENIN-1 antibody G78 and histone H3 of protein extracted from *Camerata robusta* showing higher intensity signal in piece 1 (anterior part of the body) than in piece 6 (tail area). Molecular weight marker and G78 band position as indicated. **b,** Fluorometric Western blot probed with the anti-FERRITIN antibody EO95 (green) and histone H3 (red) of protein extracted from *Cura pinguis* and *Planaria torva*. The much higher band intensity in sexually mature adults as compared to immature hatchlings supports the utility of EO59 as yolk marker in other species.

1 Egger, B., Gschwentner, R. & Rieger, R. Free-living flatworms under the knife: past and present. *Dev. Genes Evol.* **217**, 89-104, doi:10.1007/s00427-006-0120-5 (2007).

2 Grosbusch, A. L., Bertemes, P., Kauffmann, B., Gotsis, C. & Egger, B. Do not lose your head over the unequal regeneration capacity in prolecithophoran flatworms. *Biology* **11**, doi:10.3390/biology11111588 (2022).

**Supplementary tables**

| | Outgroups (Mlig and Pvit: more phylogenetically distant to planarians than Cyli and Kamp) | | | | Triclads-planarians | | | | | | | | | | | | | | | | | | | | | | | | | | | | | | | | | | | | | | | | | | | | |
|---|---|---|---|---|---|---|---|---|---|---|---|---|---|---|---|---|---|---|---|---|---|---|---|---|---|---|---|---|---|---|---|---|---|---|---|---|---|---|---|---|---|---|---|---|---|---|---|---|---|
| | | | | | Maricola | | | | | | | Cavernicola | Geoplanoidea | | | | | | | | | | | | | | | Dendrocoelidae | | | | | | | | Planariidae | | | | | | | | | | | | | |
| BUSCO genes absent from > 90% of the transcriptomes | Mlig | Pvit | Cyli | Kamp | Came | Chas | Bcan | Sdio | Ptri | Plit | Ppie | Rhod | Spas | Phaa | Gdor | Gsex | Smed | Terr | Cfor | Djap | Spab | Spat | Cura | Dsic | Slug | Dtah | Spol | Bang | Pflu | Dlac | Pcav | Dmac | Dcas | Bbru | Bann | Pnig | Roma | Pmor | Pslb | Pten | Pgra | Seid | Phpy | Hret | Bipa | Pfel | Psap | Calp | Ptoc |
| 14776at33208 | 1 | 1 | 0 | 0 | 0 | 0 | 0 | 0 | 0 | 0 | 0 | 0 | 0 | 0 | 0 | 0 | 0 | 0 | 0 | 0 | 0 | 0 | 0 | 0 | 0 | 0 | 0 | 0 | 0 | 0 | 0 | 0 | 0 | 0 | 0 | 0 | 0 | 0 | 0 | 0 | 0 | 0 | 0 | 0 | 0 | 0 | 0 | 0 | 0 |
| 65302at33208 | 1 | 1 | 0 | 0 | 0 | 0 | 0 | 0 | 0 | 0 | 0 | 0 | 0 | 0 | 0 | 0 | 0 | 0 | 0 | 0 | 0 | 0 | 0 | 0 | 0 | 0 | 0 | 0 | 0 | 0 | 0 | 0 | 0 | 0 | 0 | 0 | 0 | 0 | 0 | 0 | 0 | 0 | 0 | 0 | 0 | 0 | 0 | 0 | 0 |
| 121127at33208 | 1 | 1 | 0 | 0 | 0 | 0 | 0 | 0 | 0 | 0 | 0 | 0 | 0 | 0 | 0 | 0 | 0 | 0 | 0 | 0 | 0 | 0 | 0 | 0 | 0 | 0 | 0 | 0 | 0 | 0 | 0 | 0 | 0 | 0 | 0 | 0 | 0 | 0 | 0 | 0 | 0 | 0 | 0 | 0 | 0 | 0 | 0 | 0 | 0 |
| 152327at33208 | 1 | 1 | 0 | 0 | 0 | 0 | 0 | 0 | 0 | 0 | 0 | 0 | 0 | 0 | 0 | 0 | 0 | 0 | 0 | 0 | 0 | 0 | 0 | 0 | 0 | 0 | 0 | 0 | 0 | 0 | 0 | 0 | 0 | 0 | 0 | 0 | 0 | 0 | 0 | 0 | 0 | 0 | 0 | 0 | 0 | 0 | 0 | 0 | 0 |
| 191938at33208 | 1 | 1 | 1 | 0 | 0 | 0 | 0 | 0 | 0 | 0 | 0 | 0 | 0 | 0 | 0 | 0 | 0 | 0 | 0 | 0 | 0 | 0 | 0 | 0 | 0 | 0 | 0 | 0 | 0 | 0 | 0 | 0 | 0 | 0 | 0 | 0 | 0 | 0 | 0 | 0 | 0 | 0 | 0 | 0 | 0 | 0 | 0 | 0 | 0 |
| 205220at33208 | 1 | 1 | 0 | 0 | 1 | 0 | 0 | 0 | 0 | 0 | 1 | 0 | 0 | 0 | 0 | 0 | 0 | 0 | 0 | 0 | 0 | 0 | 0 | 0 | 0 | 0 | 0 | 0 | 0 | 0 | 0 | 0 | 0 | 0 | 0 | 0 | 0 | 0 | 0 | 0 | 0 | 0 | 0 | 0 | 0 | 0 | 0 | 0 | 0 |
| 302921at33208 | 1 | 1 | 0 | 0 | 0 | 0 | 0 | 0 | 0 | 0 | 0 | 0 | 0 | 0 | 0 | 0 | 0 | 0 | 0 | 0 | 0 | 0 | 0 | 0 | 0 | 0 | 0 | 0 | 0 | 0 | 0 | 0 | 0 | 0 | 0 | 0 | 0 | 0 | 0 | 0 | 0 | 0 | 0 | 0 | 0 | 0 | 0 | 0 | 0 |
| 336647at33208 | 1 | 1 | 0 | 0 | 0 | 0 | 0 | 0 | 0 | 0 | 0 | 0 | 0 | 0 | 0 | 0 | 0 | 0 | 0 | 0 | 0 | 0 | 0 | 0 | 0 | 0 | 0 | 0 | 0 | 0 | 0 | 0 | 0 | 0 | 0 | 0 | 0 | 0 | 0 | 0 | 0 | 0 | 0 | 0 | 0 | 0 | 0 | 0 | 0 |
| 344611at33208 | 1 | 1 | 0 | 0 | 0 | 0 | 0 | 0 | 0 | 0 | 0 | 0 | 0 | 0 | 0 | 0 | 0 | 0 | 0 | 0 | 0 | 0 | 0 | 0 | 0 | 0 | 0 | 0 | 0 | 0 | 0 | 0 | 0 | 0 | 0 | 0 | 0 | 0 | 0 | 0 | 0 | 0 | 0 | 0 | 0 | 0 | 0 | 0 | 0 |
| 359369at33208 | 1 | 1 | 0 | 0 | 0 | 0 | 0 | 0 | 0 | 0 | 0 | 0 | 0 | 0 | 0 | 0 | 0 | 0 | 0 | 0 | 0 | 0 | 0 | 0 | 0 | 0 | 0 | 0 | 0 | 0 | 0 | 0 | 0 | 0 | 0 | 0 | 0 | 0 | 0 | 0 | 0 | 0 | 0 | 0 | 0 | 0 | 0 | 0 | 0 |
| 367490at33208 | 1 | 1 | 0 | 0 | 0 | 0 | 0 | 0 | 0 | 0 | 0 | 0 | 0 | 0 | 0 | 0 | 0 | 0 | 0 | 0 | 0 | 0 | 0 | 0 | 0 | 0 | 0 | 0 | 0 | 0 | 0 | 0 | 0 | 0 | 0 | 0 | 0 | 0 | 0 | 0 | 0 | 0 | 0 | 0 | 0 | 0 | 0 | 0 | 0 |
| 368670at33208 | 1 | 1 | 0 | 0 | 0 | 0 | 0 | 0 | 0 | 0 | 0 | 0 | 0 | 0 | 0 | 0 | 0 | 0 | 0 | 0 | 0 | 0 | 0 | 0 | 0 | 0 | 0 | 0 | 0 | 0 | 0 | 0 | 0 | 0 | 0 | 0 | 0 | 0 | 0 | 0 | 0 | 0 | 0 | 0 | 0 | 0 | 0 | 0 | 0 |
| 387079at33208 | 1 | 1 | 0 | 0 | 1 | 0 | 0 | 0 | 0 | 0 | 0 | 0 | 0 | 0 | 0 | 0 | 0 | 0 | 0 | 0 | 0 | 0 | 0 | 0 | 0 | 0 | 0 | 0 | 0 | 0 | 0 | 0 | 0 | 0 | 0 | 0 | 0 | 0 | 0 | 0 | 0 | 0 | 0 | 0 | 0 | 0 | 0 | 0 | 0 |
| 393219at33208 | 1 | 1 | 0 | 0 | 0 | 0 | 0 | 0 | 0 | 0 | 0 | 0 | 0 | 0 | 0 | 0 | 0 | 0 | 0 | 0 | 0 | 0 | 0 | 0 | 0 | 0 | 0 | 0 | 0 | 0 | 0 | 0 | 0 | 0 | 0 | 0 | 0 | 0 | 0 | 0 | 0 | 0 | 0 | 0 | 0 | 0 | 0 | 0 | 0 |
| 398202at33208 | 1 | 1 | 1 | 0 | 0 | 0 | 0 | 0 | 0 | 0 | 0 | 0 | 0 | 0 | 0 | 0 | 0 | 0 | 0 | 0 | 0 | 0 | 0 | 0 | 0 | 0 | 0 | 0 | 0 | 0 | 0 | 0 | 0 | 0 | 0 | 0 | 0 | 0 | 0 | 0 | 0 | 0 | 0 | 0 | 0 | 0 | 0 | 0 | 0 |
| 409186at33208 | 1 | 1 | 0 | 0 | 0 | 0 | 0 | 0 | 0 | 0 | 0 | 0 | 0 | 0 | 0 | 0 | 0 | 0 | 0 | 0 | 0 | 0 | 0 | 0 | 0 | 0 | 0 | 0 | 0 | 0 | 0 | 0 | 0 | 0 | 0 | 0 | 0 | 0 | 0 | 0 | 0 | 0 | 0 | 0 | 0 | 0 | 0 | 0 | 0 |
| 440195at33208 | 1 | 1 | 0 | 0 | 0 | 0 | 0 | 0 | 0 | 0 | 0 | 0 | 0 | 0 | 0 | 0 | 0 | 0 | 0 | 0 | 0 | 0 | 0 | 0 | 0 | 0 | 0 | 0 | 0 | 0 | 0 | 0 | 0 | 0 | 0 | 0 | 1 | 0 | 0 | 0 | 0 | 0 | 0 | 0 | 0 | 0 | 0 | 0 | 0 |
| 445031at33208 | 1 | 1 | 0 | 0 | 0 | 0 | 0 | 0 | 0 | 0 | 0 | 0 | 0 | 0 | 0 | 0 | 0 | 0 | 0 | 0 | 0 | 0 | 0 | 0 | 0 | 0 | 0 | 0 | 0 | 0 | 0 | 0 | 0 | 0 | 0 | 0 | 1 | 0 | 0 | 0 | 0 | 0 | 0 | 0 | 0 | 0 | 0 | 0 | 0 |
| 449056at33208 | 1 | 1 | 0 | 0 | 0 | 0 | 0 | 0 | 0 | 0 | 0 | 0 | 0 | 0 | 0 | 0 | 0 | 0 | 0 | 0 | 0 | 0 | 0 | 0 | 0 | 0 | 0 | 0 | 0 | 0 | 0 | 0 | 0 | 0 | 0 | 0 | 0 | 0 | 0 | 0 | 0 | 0 | 0 | 0 | 0 | 0 | 0 | 0 | 0 |
| 453052at33208 | 1 | 1 | 0 | 0 | 0 | 0 | 0 | 0 | 0 | 0 | 0 | 0 | 0 | 0 | 0 | 0 | 0 | 0 | 0 | 0 | 0 | 0 | 0 | 0 | 0 | 0 | 0 | 0 | 0 | 0 | 0 | 0 | 0 | 0 | 0 | 0 | 0 | 0 | 0 | 1 | 0 | 0 | 0 | 0 | 0 | 0 | 0 | 0 | 0 |
| 453056at33208 | 1 | 1 | 0 | 0 | 0 | 0 | 0 | 0 | 0 | 0 | 0 | 0 | 0 | 0 | 0 | 0 | 0 | 0 | 0 | 0 | 0 | 0 | 0 | 0 | 0 | 0 | 0 | 0 | 0 | 0 | 0 | 0 | 0 | 0 | 0 | 0 | 0 | 0 | 0 | 0 | 0 | 0 | 0 | 0 | 0 | 0 | 0 | 0 | 0 |
| 465651at33208 | 1 | 1 | 0 | 0 | 0 | 0 | 0 | 0 | 0 | 0 | 0 | 0 | 0 | 0 | 0 | 0 | 0 | 0 | 0 | 0 | 0 | 0 | 0 | 0 | 0 | 0 | 0 | 0 | 0 | 0 | 0 | 0 | 0 | 0 | 0 | 0 | 0 | 0 | 0 | 0 | 0 | 0 | 0 | 0 | 0 | 0 | 0 | 0 | 0 |
| 480700at33208 | 1 | 1 | 0 | 0 | 0 | 0 | 0 | 0 | 0 | 0 | 0 | 0 | 0 | 0 | 0 | 0 | 0 | 0 | 0 | 0 | 0 | 0 | 0 | 0 | 0 | 0 | 0 | 0 | 0 | 1 | 0 | 0 | 0 | 0 | 0 | 0 | 0 | 0 | 0 | 0 | 0 | 0 | 0 | 0 | 0 | 0 | 0 | 0 | 0 |
| 510174at33208 | 1 | 1 | 0 | 0 | 0 | 0 | 0 | 0 | 0 | 0 | 0 | 0 | 0 | 0 | 0 | 0 | 0 | 0 | 0 | 0 | 0 | 0 | 0 | 0 | 0 | 0 | 0 | 0 | 0 | 0 | 0 | 0 | 0 | 0 | 0 | 0 | 0 | 0 | 0 | 0 | 0 | 0 | 0 | 1 | 0 | 0 | 0 | 0 | 0 |
| 526839at33208 | 1 | 1 | 0 | 0 | 0 | 0 | 0 | 0 | 0 | 0 | 0 | 0 | 0 | 0 | 0 | 0 | 0 | 0 | 0 | 0 | 0 | 0 | 0 | 0 | 0 | 0 | 0 | 0 | 0 | 0 | 0 | 0 | 0 | 0 | 0 | 0 | 0 | 0 | 0 | 0 | 0 | 0 | 0 | 0 | 0 | 0 | 0 | 0 | 0 |
| 548444at33208 | 1 | 1 | 0 | 1 | 0 | 0 | 0 | 0 | 0 | 0 | 0 | 0 | 0 | 0 | 0 | 0 | 0 | 0 | 0 | 0 | 0 | 0 | 0 | 0 | 0 | 0 | 0 | 0 | 0 | 0 | 0 | 0 | 0 | 0 | 0 | 0 | 0 | 0 | 0 | 0 | 0 | 0 | 0 | 0 | 0 | 0 | 0 | 0 | 0 |
| 551632at33208 | 1 | 1 | 0 | 0 | 0 | 0 | 0 | 0 | 0 | 0 | 0 | 0 | 0 | 0 | 0 | 0 | 0 | 0 | 0 | 0 | 0 | 0 | 0 | 0 | 0 | 0 | 0 | 0 | 0 | 0 | 0 | 0 | 0 | 0 | 0 | 0 | 0 | 0 | 0 | 0 | 0 | 0 | 0 | 0 | 0 | 0 | 0 | 0 | 0 |
| 552764at33208 | 1 | 1 | 0 | 0 | 0 | 0 | 0 | 0 | 0 | 0 | 0 | 1 | 0 | 0 | 0 | 0 | 0 | 0 | 0 | 0 | 0 | 0 | 0 | 0 | 0 | 0 | 0 | 0 | 0 | 0 | 0 | 0 | 0 | 0 | 0 | 0 | 1 | 0 | 0 | 0 | 0 | 0 | 0 | 0 | 0 | 0 | 0 | 0 | 0 |
| 556189at33208 | 1 | 1 | 0 | 0 | 0 | 0 | 0 | 0 | 0 | 0 | 0 | 0 | 0 | 0 | 0 | 0 | 0 | 0 | 0 | 0 | 0 | 0 | 0 | 0 | 0 | 0 | 0 | 0 | 0 | 0 | 0 | 0 | 0 | 0 | 0 | 0 | 0 | 0 | 0 | 0 | 0 | 0 | 0 | 0 | 0 | 0 | 0 | 0 | 0 |
| 556510at33208 | 1 | 1 | 0 | 0 | 0 | 0 | 0 | 0 | 0 | 0 | 0 | 0 | 0 | 0 | 0 | 0 | 0 | 0 | 0 | 0 | 0 | 0 | 0 | 0 | 0 | 0 | 0 | 0 | 0 | 0 | 0 | 0 | 0 | 0 | 0 | 0 | 0 | 0 | 0 | 0 | 0 | 0 | 0 | 0 | 0 | 0 | 0 | 0 | 0 |
| 559576at33208 | 1 | 1 | 0 | 0 | 0 | 0 | 0 | 0 | 0 | 0 | 0 | 0 | 0 | 0 | 0 | 0 | 0 | 0 | 0 | 0 | 0 | 0 | 0 | 0 | 0 | 0 | 0 | 0 | 0 | 0 | 0 | 0 | 0 | 0 | 0 | 0 | 0 | 0 | 0 | 0 | 0 | 0 | 0 | 0 | 0 | 0 | 0 | 0 | 0 |
| 569303at33208 | 1 | 1 | 0 | 0 | 0 | 0 | 0 | 0 | 0 | 0 | 0 | 0 | 0 | 1 | 0 | 0 | 0 | 0 | 0 | 0 | 0 | 0 | 0 | 0 | 0 | 0 | 0 | 0 | 0 | 0 | 0 | 0 | 0 | 0 | 0 | 0 | 0 | 0 | 0 | 0 | 0 | 0 | 0 | 0 | 0 | 0 | 0 | 0 | 0 |
| 571149at33208 | 1 | 1 | 0 | 0 | 0 | 0 | 0 | 0 | 0 | 0 | 0 | 0 | 0 | 0 | 0 | 0 | 0 | 0 | 0 | 0 | 0 | 0 | 0 | 0 | 0 | 0 | 0 | 0 | 0 | 0 | 0 | 0 | 0 | 0 | 0 | 0 | 0 | 0 | 0 | 0 | 0 | 0 | 1 | 0 | 0 | 0 | 0 | 0 | 0 |
| 572463at33208 | 1 | 1 | 0 | 0 | 0 | 0 | 0 | 0 | 0 | 0 | 0 | 0 | 0 | 0 | 0 | 0 | 0 | 0 | 0 | 0 | 0 | 0 | 0 | 0 | 0 | 0 | 0 | 0 | 0 | 0 | 0 | 0 | 0 | 0 | 0 | 0 | 1 | 0 | 0 | 0 | 0 | 0 | 0 | 0 | 0 | 0 | 0 | 0 | 0 |
| 585313at33208 | 1 | 1 | 0 | 0 | 0 | 0 | 0 | 0 | 0 | 0 | 1 | 0 | 0 | 0 | 0 | 0 | 0 | 0 | 0 | 0 | 0 | 0 | 0 | 0 | 0 | 0 | 0 | 0 | 0 | 0 | 0 | 0 | 0 | 0 | 0 | 0 | 0 | 0 | 0 | 0 | 0 | 0 | 0 | 0 | 0 | 0 | 0 | 0 | 0 |
| 608669at33208 | 1 | 1 | 1 | 0 | 0 | 0 | 0 | 0 | 0 | 0 | 0 | 0 | 0 | 0 | 0 | 0 | 0 | 0 | 0 | 0 | 0 | 0 | 0 | 1 | 0 | 0 | 0 | 0 | 0 | 0 | 0 | 0 | 0 | 0 | 0 | 0 | 0 | 0 | 0 | 0 | 0 | 0 | 0 | 0 | 0 | 0 | 0 | 0 | 0 |
| 203698at33208 | 1 | 0 | 0 | 0 | 0 | 0 | 0 | 0 | 0 | 0 | 0 | 0 | 0 | 0 | 0 | 0 | 0 | 0 | 0 | 0 | 0 | 0 | 0 | 0 | 0 | 0 | 0 | 0 | 0 | 0 | 0 | 0 | 0 | 0 | 0 | 0 | 0 | 0 | 0 | 0 | 0 | 0 | 0 | 0 | 0 | 0 | 0 | 0 | 0 |
| 304388at33208 | 1 | 0 | 0 | 0 | 0 | 0 | 0 | 0 | 0 | 0 | 0 | 0 | 0 | 0 | 0 | 0 | 0 | 0 | 0 | 0 | 0 | 0 | 0 | 0 | 0 | 0 | 0 | 0 | 0 | 0 | 0 | 0 | 0 | 0 | 0 | 0 | 0 | 0 | 0 | 0 | 0 | 0 | 0 | 0 | 0 | 0 | 0 | 0 | 0 |
| 442169at33208 | 1 | 0 | 0 | 0 | 0 | 0 | 0 | 0 | 0 | 0 | 0 | 0 | 0 | 0 | 0 | 0 | 0 | 0 | 0 | 0 | 0 | 0 | 0 | 0 | 0 | 0 | 0 | 0 | 0 | 0 | 0 | 0 | 0 | 0 | 0 | 0 | 0 | 0 | 0 | 0 | 0 | 0 | 0 | 0 | 0 | 0 | 0 | 0 | 0 |
| 489359at33208 | 1 | 0 | 0 | 0 | 0 | 0 | 0 | 0 | 0 | 0 | 0 | 0 | 0 | 0 | 0 | 0 | 0 | 0 | 0 | 0 | 0 | 0 | 0 | 0 | 0 | 0 | 0 | 0 | 0 | 1 | 0 | 0 | 0 | 0 | 0 | 0 | 0 | 0 | 0 | 0 | 0 | 0 | 0 | 0 | 0 | 0 | 0 | 0 | 0 |
| 495802at33208 | 1 | 0 | 0 | 0 | 0 | 0 | 0 | 0 | 0 | 0 | 0 | 0 | 0 | 0 | 0 | 0 | 0 | 0 | 0 | 0 | 0 | 0 | 0 | 0 | 0 | 0 | 0 | 0 | 0 | 0 | 0 | 0 | 0 | 0 | 0 | 0 | 0 | 0 | 0 | 0 | 0 | 0 | 0 | 0 | 0 | 0 | 0 | 0 | 0 |
| 544849at33208 | 1 | 0 | 0 | 0 | 0 | 0 | 0 | 0 | 0 | 0 | 0 | 0 | 0 | 0 | 0 | 0 | 0 | 0 | 0 | 0 | 0 | 0 | 0 | 0 | 0 | 0 | 0 | 0 | 0 | 0 | 0 | 0 | 0 | 0 | 0 | 0 | 0 | 0 | 0 | 0 | 0 | 0 | 0 | 0 | 0 | 0 | 0 | 0 | 0 |
| 578684at33208 | 1 | 0 | 0 | 0 | 0 | 0 | 0 | 0 | 0 | 0 | 0 | 0 | 0 | 0 | 0 | 0 | 0 | 0 | 0 | 0 | 0 | 0 | 0 | 0 | 0 | 0 | 0 | 0 | 0 | 0 | 0 | 0 | 0 | 0 | 0 | 0 | 0 | 0 | 0 | 0 | 0 | 0 | 0 | 0 | 0 | 0 | 0 | 0 | 0 |
| 591316at33208 | 1 | 0 | 1 | 1 | 0 | 0 | 0 | 0 | 0 | 0 | 0 | 0 | 0 | 0 | 0 | 0 | 0 | 0 | 0 | 0 | 0 | 0 | 0 | 0 | 0 | 0 | 0 | 0 | 0 | 0 | 0 | 0 | 0 | 0 | 0 | 0 | 0 | 0 | 0 | 0 | 0 | 0 | 0 | 0 | 0 | 0 | 0 | 0 | 0 |
| 606795at33208 | 1 | 0 | 0 | 0 | 0 | 0 | 0 | 0 | 0 | 0 | 0 | 0 | 0 | 0 | 0 | 0 | 0 | 0 | 0 | 0 | 0 | 0 | 0 | 0 | 0 | 0 | 0 | 0 | 0 | 0 | 0 | 0 | 0 | 0 | 0 | 0 | 0 | 0 | 0 | 0 | 0 | 0 | 0 | 0 | 0 | 0 | 0 | 0 | 0 |
| 628082at33208 | 1 | 0 | 0 | 0 | 0 | 0 | 0 | 0 | 0 | 0 | 0 | 0 | 0 | 0 | 1 | 1 | 0 | 0 | 0 | 0 | 0 | 0 | 0 | 0 | 0 | 0 | 1 | 0 | 0 | 0 | 0 | 0 | 0 | 0 | 0 | 0 | 0 | 0 | 0 | 0 | 0 | 0 | 0 | 0 | 0 | 0 | 0 | 0 | 0 |
| 629763at33208 | 1 | 0 | 0 | 0 | 0 | 0 | 0 | 0 | 0 | 0 | 0 | 0 | 0 | 0 | 0 | 0 | 0 | 0 | 0 | 0 | 0 | 0 | 0 | 0 | 0 | 0 | 0 | 0 | 0 | 1 | 0 | 0 | 0 | 0 | 0 | 0 | 0 | 0 | 0 | 0 | 0 | 0 | 0 | 0 | 0 | 0 | 0 | 0 | 0 |
| 157138at33208 | 0 | 1 | 0 | 0 | 0 | 0 | 0 | 0 | 0 | 0 | 0 | 0 | 0 | 0 | 0 | 0 | 0 | 0 | 0 | 0 | 0 | 0 | 0 | 0 | 0 | 0 | 0 | 0 | 0 | 0 | 0 | 0 | 0 | 0 | 0 | 0 | 1 | 0 | 0 | 0 | 0 | 0 | 0 | 0 | 0 | 0 | 0 | 0 | 0 |
| 261614at33208 | 0 | 1 | 0 | 0 | 0 | 0 | 0 | 0 | 0 | 0 | 0 | 0 | 0 | 0 | 0 | 0 | 0 | 0 | 0 | 0 | 0 | 0 | 0 | 0 | 0 | 0 | 0 | 0 | 0 | 0 | 0 | 0 | 0 | 0 | 0 | 0 | 0 | 0 | 0 | 0 | 0 | 0 | 0 | 0 | 0 | 0 | 0 | 0 | 0 |
| 331186at33208 | 0 | 1 | 0 | 0 | 0 | 0 | 0 | 0 | 0 | 0 | 0 | 0 | 0 | 0 | 0 | 0 | 0 | 0 | 0 | 0 | 0 | 0 | 0 | 0 | 0 | 0 | 0 | 0 | 0 | 0 | 0 | 0 | 0 | 0 | 0 | 0 | 0 | 0 | 0 | 0 | 0 | 0 | 0 | 0 | 0 | 0 | 0 | 0 | 0 |
| 356175at33208 | 0 | 1 | 0 | 0 | 0 | 0 | 0 | 0 | 0 | 0 | 0 | 0 | 0 | 0 | 0 | 0 | 0 | 0 | 0 | 0 | 0 | 0 | 0 | 0 | 0 | 0 | 0 | 0 | 0 | 0 | 0 | 0 | 0 | 0 | 0 | 0 | 0 | 0 | 0 | 0 | 0 | 0 | 0 | 0 | 0 | 0 | 0 | 0 | 0 |
| 359532at33208 | 0 | 1 | 0 | 0 | 0 | 0 | 0 | 0 | 0 | 0 | 0 | 0 | 0 | 0 | 0 | 0 | 0 | 0 | 0 | 0 | 0 | 0 | 0 | 0 | 0 | 0 | 0 | 0 | 0 | 0 | 0 | 0 | 0 | 0 | 0 | 0 | 0 | 0 | 0 | 0 | 0 | 0 | 0 | 0 | 0 | 0 | 0 | 0 | 0 |
| 365323at33208 | 0 | 1 | 0 | 0 | 0 | 0 | 0 | 0 | 0 | 0 | 0 | 0 | 0 | 0 | 0 | 0 | 0 | 0 | 0 | 0 | 0 | 0 | 0 | 0 | 0 | 0 | 0 | 0 | 0 | 0 | 0 | 0 | 0 | 0 | 0 | 0 | 0 | 0 | 0 | 0 | 0 | 0 | 0 | 0 | 0 | 0 | 0 | 0 | 0 |
| 403171at33208 | 0 | 1 | 0 | 0 | 0 | 0 | 0 | 0 | 0 | 0 | 0 | 0 | 0 | 0 | 0 | 0 | 0 | 0 | 0 | 0 | 0 | 0 | 0 | 0 | 0 | 0 | 0 | 0 | 0 | 0 | 0 | 0 | 0 | 0 | 0 | 0 | 0 | 0 | 0 | 0 | 0 | 0 | 0 | 0 | 0 | 0 | 0 | 0 | 0 |
| 430701at33208 | 0 | 1 | 0 | 0 | 0 | 0 | 0 | 0 | 0 | 0 | 0 | 0 | 0 | 0 | 0 | 0 | 0 | 0 | 0 | 0 | 1 | 0 | 0 | 0 | 0 | 0 | 0 | 0 | 0 | 0 | 0 | 0 | 0 | 0 | 0 | 0 | 0 | 0 | 0 | 0 | 0 | 0 | 0 | 0 | 0 | 0 | 0 | 0 | 0 |
| 450643at33208 | 0 | 1 | 0 | 0 | 1 | 0 | 0 | 0 | 0 | 0 | 0 | 0 | 0 | 0 | 0 | 1 | 0 | 0 | 0 | 0 | 0 | 0 | 0 | 0 | 0 | 0 | 0 | 0 | 0 | 1 | 0 | 0 | 0 | 0 | 0 | 0 | 0 | 0 | 0 | 0 | 0 | 0 | 0 | 0 | 0 | 0 | 0 | 0 | 0 |
| 463071at33208 | 0 | 1 | 0 | 0 | 0 | 0 | 0 | 0 | 0 | 0 | 0 | 0 | 0 | 0 | 0 | 0 | 0 | 0 | 0 | 0 | 0 | 0 | 0 | 0 | 0 | 0 | 0 | 0 | 0 | 0 | 0 | 0 | 0 | 0 | 0 | 0 | 0 | 0 | 0 | 0 | 0 | 0 | 0 | 0 | 0 | 0 | 0 | 0 | 0 |
| 464243at33208 | 0 | 1 | 0 | 0 | 0 | 0 | 0 | 0 | 0 | 0 | 0 | 0 | 0 | 0 | 0 | 0 | 0 | 0 | 0 | 0 | 0 | 0 | 0 | 0 | 0 | 0 | 0 | 0 | 0 | 0 | 0 | 0 | 0 | 0 | 0 | 0 | 0 | 0 | 0 | 0 | 0 | 0 | 0 | 0 | 0 | 0 | 0 | 0 | 0 |
| 465401at33208 | 0 | 1 | 0 | 0 | 0 | 0 | 0 | 0 | 0 | 0 | 0 | 0 | 0 | 0 | 0 | 0 | 0 | 0 | 0 | 0 | 0 | 0 | 0 | 0 | 0 | 0 | 0 | 0 | 0 | 0 | 0 | 0 | 0 | 0 | 0 | 0 | 0 | 0 | 0 | 0 | 0 | 0 | 0 | 0 | 0 | 0 | 0 | 0 | 0 |
| 477052at33208 | 0 | 1 | 0 | 0 | 0 | 0 | 0 | 0 | 0 | 0 | 0 | 0 | 0 | 0 | 0 | 0 | 0 | 0 | 0 | 0 | 0 | 0 | 0 | 0 | 0 | 0 | 0 | 0 | 0 | 0 | 0 | 0 | 0 | 0 | 0 | 0 | 0 | 0 | 0 | 0 | 0 | 0 | 0 | 0 | 0 | 0 | 0 | 0 | 0 |
| 493505at33208 | 0 | 1 | 0 | 0 | 0 | 0 | 0 | 0 | 0 | 0 | 0 | 0 | 0 | 0 | 0 | 0 | 0 | 0 | 0 | 0 | 0 | 0 | 0 | 0 | 0 | 0 | 0 | 0 | 0 | 1 | 0 | 0 | 0 | 0 | 0 | 0 | 0 | 0 | 0 | 0 | 0 | 0 | 0 | 0 | 0 | 0 | 0 | 0 | 0 |
| 521343at33208 | 0 | 1 | 0 | 1 | 0 | 0 | 0 | 0 | 0 | 0 | 0 | 0 | 0 | 0 | 0 | 0 | 0 | 0 | 0 | 0 | 0 | 0 | 0 | 0 | 0 | 0 | 0 | 0 | 0 | 0 | 0 | 0 | 0 | 0 | 0 | 0 | 0 | 0 | 0 | 0 | 0 | 0 | 0 | 0 | 0 | 0 | 0 | 0 | 0 |
| 535782at33208 | 0 | 1 | 0 | 0 | 0 | 0 | 0 | 0 | 0 | 0 | 0 | 0 | 0 | 0 | 0 | 0 | 0 | 0 | 0 | 0 | 0 | 0 | 0 | 0 | 0 | 0 | 0 | 0 | 0 | 0 | 0 | 0 | 0 | 0 | 0 | 0 | 0 | 0 | 0 | 0 | 0 | 0 | 0 | 0 | 0 | 0 | 0 | 0 | 0 |
| 547099at33208 | 0 | 1 | 0 | 0 | 0 | 0 | 0 | 0 | 0 | 0 | 0 | 0 | 0 | 0 | 0 | 0 | 0 | 0 | 0 | 0 | 0 | 0 | 0 | 0 | 0 | 0 | 0 | 0 | 0 | 0 | 0 | 0 | 0 | 0 | 0 | 0 | 0 | 0 | 0 | 0 | 0 | 0 | 0 | 0 | 0 | 0 | 0 | 0 | 0 |
| 549988at33208 | 0 | 1 | 1 | 0 | 0 | 0 | 0 | 0 | 0 | 0 | 0 | 0 | 0 | 0 | 0 | 0 | 0 | 0 | 0 | 0 | 0 | 0 | 0 | 0 | 0 | 0 | 0 | 0 | 0 | 0 | 0 | 0 | 0 | 0 | 0 | 0 | 0 | 0 | 0 | 0 | 0 | 0 | 0 | 0 | 0 | 0 | 0 | 0 | 0 |
| 559146at33208 | 0 | 1 | 0 | 0 | 0 | 0 | 0 | 0 | 0 | 0 | 0 | 0 | 0 | 0 | 0 | 0 | 0 | 0 | 0 | 0 | 0 | 0 | 0 | 0 | 0 | 0 | 0 | 0 | 0 | 1 | 0 | 0 | 0 | 0 | 0 | 0 | 0 | 0 | 0 | 0 | 0 | 0 | 1 | 0 | 0 | 0 | 0 | 0 | 0 |
| 581717at33208 | 0 | 1 | 0 | 0 | 0 | 0 | 0 | 0 | 0 | 0 | 0 | 0 | 0 | 0 | 0 | 0 | 0 | 0 | 0 | 0 | 0 | 0 | 0 | 0 | 0 | 0 | 0 | 0 | 0 | 0 | 0 | 0 | 0 | 0 | 0 | 0 | 0 | 0 | 0 | 0 | 0 | 0 | 0 | 0 | 0 | 0 | 0 | 0 | 0 |
| 582951at33208 | 0 | 1 | 0 | 0 | 0 | 0 | 0 | 0 | 0 | 0 | 0 | 0 | 0 | 0 | 0 | 0 | 0 | 0 | 0 | 0 | 0 | 0 | 0 | 0 | 0 | 0 | 0 | 0 | 0 | 0 | 0 | 0 | 0 | 0 | 0 | 0 | 0 | 0 | 0 | 0 | 0 | 0 | 0 | 0 | 0 | 0 | 0 | 0 | 0 |
| 590221at33208 | 0 | 1 | 0 | 0 | 0 | 0 | 0 | 0 | 0 | 0 | 0 | 0 | 0 | 0 | 0 | 0 | 0 | 0 | 0 | 0 | 0 | 0 | 0 | 0 | 0 | 0 | 0 | 0 | 0 | 0 | 0 | 0 | 0 | 0 | 0 | 0 | 0 | 0 | 0 | 0 | 0 | 0 | 0 | 0 | 0 | 0 | 0 | 0 | 0 |
| 601875at33208 | 0 | 1 | 0 | 0 | 0 | 0 | 0 | 0 | 0 | 0 | 0 | 0 | 0 | 0 | 0 | 0 | 0 | 0 | 0 | 0 | 0 | 0 | 0 | 0 | 0 | 0 | 0 | 0 | 0 | 0 | 1 | 0 | 1 | 0 | 0 | 0 | 0 | 0 | 1 | 0 | 0 | 0 | 0 | 0 | 0 | 0 | 0 | 0 | 0 |
| 602587at33208 | 0 | 1 | 0 | 0 | 0 | 0 | 0 | 0 | 0 | 0 | 0 | 0 | 0 | 0 | 0 | 0 | 0 | 0 | 0 | 0 | 0 | 0 | 0 | 0 | 0 | 0 | 0 | 0 | 0 | 0 | 0 | 0 | 0 | 0 | 0 | 0 | 0 | 0 | 0 | 0 | 0 | 0 | 0 | 0 | 0 | 0 | 0 | 0 | 0 |
| 606958at33208 | 0 | 1 | 0 | 0 | 0 | 0 | 0 | 0 | 0 | 0 | 0 | 0 | 0 | 0 | 0 | 0 | 0 | 0 | 0 | 0 | 0 | 0 | 0 | 0 | 0 | 0 | 0 | 0 | 0 | 0 | 0 | 0 | 0 | 0 | 0 | 0 | 0 | 0 | 0 | 0 | 0 | 0 | 0 | 0 | 0 | 0 | 0 | 0 | 0 |
| 623995at33208 | 0 | 1 | 0 | 0 | 0 | 0 | 0 | 0 | 0 | 0 | 0 | 0 | 0 | 0 | 1 | 0 | 0 | 0 | 0 | 0 | 0 | 0 | 0 | 0 | 0 | 0 | 0 | 0 | 0 | 0 | 0 | 0 | 0 | 0 | 0 | 0 | 0 | 0 | 0 | 0 | 0 | 0 | 0 | 0 | 0 | 0 | 0 | 0 | 0 |
| 633007at33208 | 0 | 1 | 0 | 0 | 0 | 0 | 0 | 0 | 0 | 0 | 0 | 0 | 0 | 0 | 0 | 0 | 0 | 0 | 0 | 0 | 0 | 0 | 0 | 0 | 0 | 0 | 0 | 0 | 0 | 0 | 0 | 0 | 0 | 0 | 0 | 0 | 0 | 0 | 0 | 0 | 0 | 0 | 0 | 0 | 0 | 0 | 0 | 0 | 0 |
| 502584at33208 | 0 | 0 | 0 | 0 | 0 | 0 | 0 | 0 | 0 | 0 | 0 | 0 | 0 | 0 | 0 | 0 | 0 | 0 | 0 | 0 | 0 | 0 | 0 | 0 | 0 | 0 | 0 | 0 | 0 | 1 | 0 | 0 | 0 | 0 | 0 | 0 | 0 | 0 | 0 | 0 | 0 | 0 | 0 | 0 | 0 | 0 | 0 | 0 | 0 |
| 515377at33208 | 0 | 0 | 0 | 0 | 0 | 0 | 0 | 1 | 0 | 0 | 0 | 0 | 0 | 0 | 0 | 0 | 0 | 0 | 0 | 0 | 0 | 0 | 0 | 0 | 0 | 0 | 0 | 0 | 0 | 0 | 0 | 0 | 0 | 0 | 0 | 0 | 0 | 0 | 0 | 0 | 0 | 1 | 0 | 0 | 0 | 0 | 0 | 0 | 0 |
| 558368at33208 | 0 | 0 | 0 | 1 | 0 | 0 | 0 | 0 | 0 | 0 | 0 | 0 | 0 | 0 | 0 | 0 | 0 | 0 | 0 | 0 | 0 | 0 | 0 | 0 | 0 | 0 | 0 | 0 | 0 | 0 | 0 | 0 | 0 | 0 | 0 | 0 | 0 | 0 | 0 | 0 | 0 | 0 | 0 | 0 | 0 | 0 | 0 | 0 | 0 |
| 560566at33208 | 0 | 0 | 0 | 0 | 0 | 0 | 0 | 0 | 0 | 0 | 0 | 0 | 0 | 0 | 0 | 0 | 0 | 0 | 0 | 0 | 0 | 0 | 0 | 0 | 0 | 0 | 0 | 0 | 0 | 1 | 0 | 0 | 0 | 0 | 0 | 0 | 0 | 0 | 0 | 0 | 0 | 0 | 0 | 0 | 0 | 0 | 0 | 0 | 0 |
| 590470at33208 | 0 | 0 | 0 | 0 | 0 | 0 | 0 | 0 | 0 | 0 | 0 | 0 | 0 | 0 | 0 | 0 | 0 | 0 | 0 | 0 | 0 | 0 | 0 | 0 | 0 | 0 | 0 | 0 | 0 | 0 | 0 | 0 | 0 | 0 | 0 | 0 | 0 | 0 | 0 | 0 | 0 | 0 | 1 | 0 | 0 | 0 | 0 | 0 | 0 |
| 92367at33208 | 0 | 0 | 0 | 0 | 0 | 0 | 0 | 0 | 0 | 0 | 0 | 0 | 0 | 0 | 0 | 0 | 0 | 0 | 0 | 0 | 0 | 0 | 0 | 0 | 0 | 0 | 0 | 0 | 0 | 0 | 0 | 0 | 0 | 0 | 0 | 0 | 0 | 0 | 0 | 0 | 0 | 0 | 0 | 0 | 0 | 0 | 0 | 0 | 0 |
| 108764at33208 | 0 | 0 | 0 | 0 | 0 | 0 | 0 | 0 | 0 | 0 | 0 | 0 | 0 | 0 | 0 | 0 | 0 | 0 | 0 | 0 | 0 | 0 | 0 | 0 | 0 | 0 | 0 | 0 | 0 | 0 | 0 | 0 | 0 | 0 | 0 | 0 | 0 | 0 | 0 | 0 | 0 | 0 | 0 | 0 | 0 | 0 | 0 | 0 | 0 |
| 111746at33208 | 0 | 0 | 0 | 0 | 0 | 0 | 0 | 0 | 0 | 0 | 0 | 0 | 0 | 0 | 0 | 0 | 0 | 0 | 0 | 0 | 0 | 0 | 0 | 0 | 0 | 0 | 0 | 0 | 0 | 0 | 0 | 0 | 0 | 0 | 0 | 0 | 0 | 0 | 0 | 0 | 0 | 0 | 0 | 0 | 0 | 0 | 0 | 0 | 0 |
| 216257at33208 | 0 | 0 | 0 | 0 | 0 | 0 | 0 | 0 | 0 | 0 | 0 | 0 | 0 | 0 | 0 | 0 | 0 | 0 | 0 | 0 | 0 | 0 | 0 | 0 | 0 | 0 | 0 | 0 | 0 | 0 | 0 | 0 | 0 | 0 | 0 | 0 | 0 | 0 | 0 | 0 | 0 | 0 | 0 | 0 | 0 | 0 | 0 | 0 | 0 |
| 239149at33208 | 0 | 0 | 0 | 0 | 0 | 0 | 0 | 0 | 0 | 0 | 0 | 0 | 0 | 0 | 0 | 0 | 0 | 0 | 0 | 0 | 0 | 0 | 0 | 0 | 0 | 0 | 0 | 0 | 0 | 0 | 0 | 0 | 0 | 0 | 0 | 0 | 0 | 0 | 0 | 0 | 0 | 0 | 0 | 0 | 0 | 0 | 0 | 0 | 0 |
| 239597at33208 | 0 | 0 | 0 | 0 | 0 | 0 | 0 | 0 | 0 | 0 | 0 | 0 | 0 | 0 | 0 | 0 | 0 | 0 | 0 | 0 | 0 | 0 | 0 | 0 | 0 | 0 | 0 | 0 | 0 | 0 | 0 | 0 | 0 | 0 | 0 | 0 | 0 | 0 | 0 | 0 | 0 | 0 | 0 | 0 | 0 | 0 | 0 | 0 | 0 |
| 272605at33208 | 0 | 0 | 0 | 0 | 0 | 0 | 0 | 0 | 0 | 0 | 0 | 0 | 0 | 0 | 0 | 0 | 0 | 0 | 0 | 0 | 0 | 0 | 0 | 0 | 0 | 0 | 0 | 0 | 0 | 0 | 0 | 0 | 0 | 0 | 0 | 0 | 0 | 0 | 0 | 0 | 0 | 0 | 0 | 0 | 0 | 0 | 0 | 0 | 0 |
| 295556at33208 | 0 | 0 | 0 | 0 | 0 | 0 | 0 | 0 | 0 | 0 | 0 | 0 | 0 | 0 | 0 | 0 | 0 | 0 | 0 | 0 | 0 | 0 | 0 | 0 | 0 | 0 | 0 | 0 | 0 | 0 | 0 | 0 | 0 | 0 | 0 | 0 | 0 | 0 | 0 | 0 | 0 | 0 | 0 | 0 | 0 | 0 | 0 | 0 | 0 |
| 311438at33208 | 0 | 0 | 0 | 0 | 0 | 0 | 0 | 0 | 0 | 0 | 0 | 0 | 0 | 0 | 0 | 0 | 0 | 0 | 0 | 0 | 0 | 0 | 0 | 0 | 0 | 0 | 0 | 0 | 0 | 0 | 0 | 0 | 0 | 0 | 0 | 0 | 0 | 0 | 0 | 0 | 0 | 0 | 0 | 0 | 0 | 0 | 0 | 0 | 0 |
| 360379at33208 | 0 | 0 | 0 | 0 | 0 | 0 | 0 | 0 | 0 | 0 | 0 | 0 | 0 | 0 | 0 | 0 | 0 | 0 | 0 | 0 | 0 | 0 | 0 | 0 | 0 | 0 | 0 | 0 | 0 | 0 | 0 | 0 | 0 | 0 | 0 | 0 | 0 | 0 | 0 | 0 | 0 | 0 | 0 | 0 | 0 | 0 | 0 | 0 | 0 |
| 361216at33208 | 0 | 0 | 0 | 0 | 0 | 0 | 0 | 0 | 0 | 0 | 0 | 0 | 0 | 0 | 0 | 0 | 0 | 0 | 0 | 0 | 0 | 0 | 0 | 0 | 0 | 0 | 0 | 0 | 0 | 0 | 0 | 0 | 0 | 0 | 0 | 0 | 0 | 0 | 0 | 0 | 0 | 0 | 0 | 0 | 0 | 0 | 0 | 0 | 0 |
| 396237at33208 | 0 | 0 | 0 | 0 | 0 | 0 | 0 | 0 | 0 | 0 | 0 | 0 | 0 | 0 | 0 | 0 | 0 | 0 | 0 | 0 | 0 | 0 | 0 | 0 | 0 | 0 | 0 | 0 | 0 | 0 | 0 | 0 | 0 | 0 | 0 | 0 | 0 | 0 | 0 | 0 | 0 | 0 | 0 | 0 | 0 | 0 | 0 | 0 | 0 |
| 399439at33208 | 0 | 0 | 0 | 0 | 0 | 0 | 0 | 0 | 0 | 0 | 0 | 0 | 0 | 0 | 0 | 0 | 0 | 0 | 0 | 0 | 0 | 0 | 0 | 0 | 0 | 0 | 0 | 0 | 0 | 0 | 0 | 0 | 0 | 0 | 0 | 0 | 0 | 0 | 0 | 0 | 0 | 0 | 0 | 0 | 0 | 0 | 0 | 0 | 0 |
| 428754at33208 | 0 | 0 | 0 | 0 | 0 | 0 | 0 | 0 | 0 | 0 | 0 | 0 | 0 | 0 | 0 | 0 | 0 | 0 | 0 | 0 | 0 | 0 | 0 | 0 | 0 | 0 | 0 | 0 | 0 | 0 | 0 | 0 | 0 | 0 | 0 | 0 | 0 | 0 | 0 | 0 | 0 | 0 | 0 | 0 | 0 | 0 | 0 | 0 | 0 |
| 435253at33208 | 0 | 0 | 0 | 0 | 0 | 0 | 0 | 0 | 0 | 0 | 0 | 0 | 0 | 0 | 0 | 0 | 0 | 0 | 0 | 0 | 0 | 0 | 0 | 0 | 0 | 0 | 0 | 0 | 0 | 0 | 0 | 0 | 0 | 0 | 0 | 0 | 0 | 0 | 0 | 0 | 0 | 0 | 0 | 0 | 0 | 0 | 0 | 0 | 0 |
| 446514at33208 | 0 | 0 | 0 | 0 | 0 | 0 | 0 | 0 | 0 | 0 | 0 | 0 | 0 | 0 | 0 | 0 | 0 | 0 | 0 | 0 | 0 | 0 | 0 | 0 | 0 | 0 | 0 | 0 | 0 | 0 | 0 | 0 | 0 | 0 | 0 | 0 | 0 | 0 | 0 | 0 | 0 | 0 | 0 | 0 | 0 | 0 | 0 | 0 | 0 |
| 456998at33208 | 0 | 0 | 0 | 0 | 0 | 0 | 0 | 0 | 0 | 0 | 0 | 0 | 0 | 0 | 0 | 0 | 0 | 0 | 0 | 0 | 0 | 0 | 0 | 0 | 0 | 0 | 0 | 0 | 0 | 0 | 0 | 0 | 0 | 0 | 0 | 0 | 0 | 0 | 0 | 0 | 0 | 0 | 0 | 0 | 0 | 0 | 0 | 0 | 0 |
| 470197at33208 | 0 | 0 | 0 | 0 | 0 | 0 | 0 | 0 | 0 | 0 | 0 | 0 | 0 | 0 | 0 | 0 | 0 | 0 | 0 | 0 | 0 | 0 | 0 | 0 | 0 | 0 | 0 | 0 | 0 | 0 | 0 | 0 | 0 | 0 | 0 | 0 | 0 | 0 | 0 | 0 | 0 | 0 | 0 | 0 | 0 | 0 | 0 | 0 | 0 |
| 474302at33208 | 0 | 0 | 0 | 0 | 0 | 0 | 0 | 0 | 0 | 0 | 0 | 0 | 0 | 0 | 0 | 0 | 0 | 0 | 0 | 0 | 0 | 0 | 0 | 0 | 0 | 0 | 0 | 0 | 0 | 0 | 0 | 0 | 0 | 0 | 0 | 0 | 0 | 0 | 0 | 0 | 0 | 0 | 0 | 0 | 0 | 0 | 0 | 0 | 0 |
| 490961at33208 | 0 | 0 | 0 | 0 | 0 | 0 | 0 | 0 | 0 | 0 | 0 | 0 | 0 | 0 | 0 | 0 | 0 | 0 | 0 | 0 | 0 | 0 | 0 | 0 | 0 | 0 | 0 | 0 | 0 | 0 | 0 | 0 | 0 | 0 | 0 | 0 | 0 | 0 | 0 | 0 | 0 | 0 | 0 | 0 | 0 | 0 | 0 | 0 | 0 |
| 495329at33208 | 0 | 0 | 0 | 0 | 0 | 0 | 0 | 0 | 0 | 0 | 0 | 0 | 0 | 0 | 0 | 0 | 0 | 0 | 0 | 0 | 0 | 0 | 0 | 0 | 0 | 0 | 0 | 0 | 0 | 0 | 0 | 0 | 0 | 0 | 0 | 0 | 0 | 0 | 0 | 0 | 0 | 0 | 0 | 0 | 0 | 0 | 0 | 0 | 0 |
| 515528at33208 | 0 | 0 | 0 | 0 | 0 | 0 | 0 | 0 | 0 | 0 | 0 | 0 | 0 | 0 | 0 | 0 | 0 | 0 | 0 | 0 | 0 | 0 | 0 | 0 | 0 | 0 | 0 | 0 | 0 | 0 | 0 | 0 | 0 | 0 | 0 | 0 | 0 | 0 | 0 | 0 | 0 | 0 | 0 | 0 | 0 | 0 | 0 | 0 | 0 |
| 534757at33208 | 0 | 0 | 0 | 0 | 0 | 0 | 0 | 0 | 0 | 0 | 0 | 0 | 0 | 0 | 0 | 0 | 0 | 0 | 0 | 0 | 0 | 0 | 0 | 0 | 0 | 0 | 0 | 0 | 0 | 0 | 0 | 0 | 0 | 0 | 0 | 0 | 0 | 0 | 0 | 0 | 0 | 0 | 0 | 0 | 0 | 0 | 0 | 0 | 0 |
| 544919at33208 | 0 | 0 | 0 | 0 | 0 | 0 | 0 | 0 | 0 | 0 | 0 | 0 | 0 | 0 | 0 | 0 | 0 | 0 | 0 | 0 | 0 | 0 | 0 | 0 | 0 | 0 | 0 | 0 | 0 | 0 | 0 | 0 | 0 | 0 | 0 | 0 | 0 | 0 | 0 | 0 | 0 | 0 | 0 | 0 | 0 | 0 | 0 | 0 | 0 |
| 558365at33208 | 0 | 0 | 0 | 0 | 0 | 0 | 0 | 0 | 0 | 0 | 0 | 0 | 0 | 0 | 0 | 0 | 0 | 0 | 0 | 0 | 0 | 0 | 0 | 0 | 0 | 0 | 0 | 0 | 0 | 0 | 0 | 0 | 0 | 0 | 0 | 0 | 0 | 0 | 0 | 0 | 0 | 0 | 0 | 0 | 0 | 0 | 0 | 0 | 0 |
| 566170at33208 | 0 | 0 | 0 | 0 | 0 | 0 | 0 | 0 | 0 | 0 | 0 | 0 | 0 | 0 | 0 | 0 | 0 | 0 | 0 | 0 | 0 | 0 | 0 | 0 | 0 | 0 | 0 | 0 | 0 | 0 | 0 | 0 | 0 | 0 | 0 | 0 | 0 | 0 | 0 | 0 | 0 | 0 | 0 | 0 | 0 | 0 | 0 | 0 | 0 |
| 613522at33208 | 0 | 0 | 0 | 0 | 0 | 0 | 0 | 0 | 0 | 0 | 0 | 0 | 0 | 0 | 0 | 0 | 0 | 0 | 0 | 0 | 0 | 0 | 0 | 0 | 0 | 0 | 0 | 0 | 0 | 0 | 0 | 0 | 0 | 0 | 0 | 0 | 0 | 0 | 0 | 0 | 0 | 0 | 0 | 0 | 0 | 0 | 0 | 0 | 0 |
| 619250at33208 | 0 | 0 | 0 | 0 | 0 | 0 | 0 | 0 | 0 | 0 | 0 | 0 | 0 | 0 | 0 | 0 | 0 | 0 | 0 | 0 | 0 | 0 | 0 | 0 | 0 | 0 | 0 | 0 | 0 | 0 | 0 | 0 | 0 | 0 | 0 | 0 | 0 | 0 | 0 | 0 | 0 | 0 | 0 | 0 | 0 | 0 | 0 | 0 | 0 |
| 638541at33208 | 0 | 0 | 0 | 0 | 0 | 0 | 0 | 0 | 0 | 0 | 0 | 0 | 0 | 0 | 0 | 0 | 0 | 0 | 0 | 0 | 0 | 0 | 0 | 0 | 0 | 0 | 0 | 0 | 0 | 0 | 0 | 0 | 0 | 0 | 0 | 0 | 0 | 0 | 0 | 0 | 0 | 0 | 0 | 0 | 0 | 0 | 0 | 0 | 0 |
| 640187at33208 | 0 | 0 | 0 | 0 | 0 | 0 | 0 | 0 | 0 | 0 | 0 | 0 | 0 | 0 | 0 | 0 | 0 | 0 | 0 | 0 | 0 | 0 | 0 | 0 | 0 | 0 | 0 | 0 | 0 | 0 | 0 | 0 | 0 | 0 | 0 | 0 | 0 | 0 | 0 | 0 | 0 | 0 | 0 | 0 | 0 | 0 | 0 | 0 | 0 |
| Number of BUSCO genes present in each transcriptome | 47 | 63 | 5 | 4 | 3 | 0 | 0 | 1 | 0 | 0 | 2 | 1 | 0 | 1 | 2 | 2 | 0 | 0 | 0 | 0 | 1 | 0 | 0 | 1 | 0 | 0 | 1 | 0 | 0 | 8 | 1 | 0 | 1 | 0 | 0 | 0 | 5 | 0 | 1 | 1 | 0 | 1 | 3 | 1 | 0 | 0 | 0 | 0 | 0 |

| | |
|---|---|
| Number of BUSCO genes absent from > 90% of the transcriptomes | 109 |
| Number of BUSCO genes absent in all transcriptomes | 30 |
| Number of BUSCO genes present in Mlig or Pvit or both, and sporadic in Cyli, Kamp and planarian transcriptomes | 74 |
| Number of BUSCO genes present in Mlig and Pvit, and sporadic in Cyli, Kamp and planarian transcriptomes | 36 |

**Supplementary Table. 1.** Presence ("1", green cell) or absence ("0", white cell) matrix of individual BUSCO genes (Y-axis) in the flatworm transcriptomes (X-axis, in phylogenetic order) (red frame in Fig. 3c), absent from > 90% of the transcriptomes. Red cells: BUSCO genes absent from all transcriptomes. Values for *S. mediterranea* are highlighted in blue for reference.

| | *dfs* | Log.lik | *AICc* | *AIC* | *⊿AICc* | *AICc*weight |
|---|---|---|---|---|---|---|
| **Dataset A** | | | | | | |
| ER | 1 | -25.8 | 53.6 | 53.7 | 2.0 | 0.3 |
| SYM | 3 | -22.3 | 51.6 | 50.7 | 0.0 | 0.7 |
| ARD | 6 | -22.0 | 59.5 | 56.0 | 7.9 | 0.0 |
| **Dataset B** | | | | | | |
| ER | 1 | -28.9 | 59.9 | 60.0 | 2.6 | 0.2 |
| SYM | 3 | -25.2 | 57.2 | 56.4 | 0.0 | 0.8 |
| ARD | 6 | -24.2 | 63.9 | 60.4 | 6.6 | 0.0 |

**Supplementary Table 2.** Model comparison for species datasets A and B. To control for the species with population-dependent variation in head regeneration ability, *Girardia dorotocephala* and *Polycelis sapporo*, we created two species datasets. Those two species are categorised in group A in dataset A and in group B in dataset B. The symmetric model is the preferred model for both datasets.

**Dataset A**

| Model | Good > Medium | Medium > Good | Medium > Bad | Bad > Medium | Good > Bad | Bad > Good | Total |
|---|---|---|---|---|---|---|---|
| ER | 5.3 | 5.4 | 4.4 | 3.3 | 3.9 | 2.8 | 25.3 |
| SYM | 4.5 | 5.9 | 0.6 | 2.2 | 0.0 | 0.0 | 13.2 |
| ARD | 4.5 | 5.8 | 0.0 | 2.1 | 0.0 | 0.0 | 12.4 |
| **Dataset B** | | | | | | | |
| ER | 4.9 | 6.3 | 4.7 | 3.4 | 3.8 | 2.6 | 25.8 |
| SYM | 5.2 | 6.3 | 0.5 | 2.2 | 0.0 | 0.0 | 14.2 |
| ARD | 6.7 | 7.1 | 0.0 | 2.1 | 0.0 | 0.0 | 15.8 |

**Supplementary Table 3.** Summaries of the number of transitions in stochastic character mappings of datasets A and B (see methods for details on dataset characteristics). Given is the average number of transitions between states across 10.000 simulated character histories. The symmetric (SYM) model was preferred in both datasets (see S. Table 2) and is highlighted in grey. Data is used in Fig. 3e and S. Fig. 3c. The number of transitions is higher than those plotted in Fig. 3e and S. Fig. 3c figures since intra-branch transitions computed in the table are not plotted in the figure.

| gene name | species | contig name | forward primer sequence (no tails) | reverse primer sequence (no tails) |
|---|---|---|---|---|
| *Spas-ß-catenin-1* | *Spathula sp.* | dd_Spas_v1_35799_1_1 | ATGATGAACGAAAGTATGAATATTGCCAATTCC | CATCGAAAATCCAAGTAGCTTGACTAAGC |
| *Smed-ß-catenin-1* | *Schmidtea mediterranea* | dd_Smed_v6_2688_0_1 | ATGATGAACGAAAGTATGAATATC | TTCCATTGAAAAACCGAGCAAT |
| *Ptor-ß-catenin-1* | *Planaria torva* | dd_Ptor_v3_7898_1_1 or dd_Ptor_v1_26557_0_1 | ATGAATGAACAGATGAATATCGTA | TTCAATAGTAAATCCAAGAAGTTTGAC |
| *Pten-ß-catenin-1* | *Polycelis tenuis* | dd_Pten_v1_68513_0_1 | ATGAATGAGCAGATGAATATGGTA | TTCCATAGAAAATCCCAGAAGTTTCACCAA |
| *Phpy-ß-catenin-1* | *Phagocata pyrenaica* | dd_Phpy_v1_51770_1_1 | ATGAATGAACACATGAATGTTGTTAATTCA | TTCCATGGTAAATCCAAGAAGTTTGAC |
| *Pfel-ß-catenin-1* | *Polycelis felina* | dd_Pfel_v1_83298_0_3 | GTTATGAACGAACAAATGAATATCG | TTCCATCGAAAATCCAAGAAGTTTAACC |
| *Gtig-ß-catenin-1* | *Girardia tigrina* | dd_Gsex_v1_24175_1_1 | ATGATGAACGAAAGTATGAATATTGTAAATTCTCC | CATTGAAAATCCCAACAATTTAACCAAACACG |
| *Dtah-ß-catenin-1* | *Dugesia tahitiense* | dd_Dtah_v1_34254_0_2 | ATGATGAACGAAAGTATGAATATTGTC | TTCCATAGAAAATCCAAGCAATTTTAC |
| *Dlac-ß-catenin-1* | *Dendrocoelum lacteum* | dd_Dlac_v8_183513_0_1 | ATGAATGAACAATTAAATTCCATGAATTCC | TTCCATAGAAAATCCAAGTAATTTTACC |
| *Djap-ß-catenin-1* | *Dugesia japonica* | dd_Djap_v1_146010_0_1 | ATGATGAACGAAAGTATGAATATTGTC | TTCCATGGAAAATCCAAGTAATTTTAC |
| *Cura-ß-catenin-1* | *Cura pinguis* | dd_Cura_v1_053333_01_01 | ATGATGAACGAAAGTATGAATATTGTTAATTC | TTCCATAGAAAATCCAAGTAATTTTACC |
| *Came-ß-catenin-1* | *Camerata robusta* | dd_Came_v1_29389_1_1 | ATGAATGACAATTCCAAGATGATAAG | TTCCATTGTAAATCCAAGCAGTTTTAC |
| *Bcan-ß-catenin-1* | *Bdelloura candida* | dd_Bcan_v1_027080_01_01 | ACAATGAATGATAATTGTAAAATGATTAATTCTCC | CTCCATTGAGAAACCAAGTAGTTTAACC |
| *Bcan-APC* | *Bdelloura candida* | dd_Bcan_v1_038064_01_08 | GCTCTGTTGGTTCATTGGACAG | TTTGGAGGTTTTGGAGGCCG |
| *Came-APC* | *Camerata robusta* | dd_Came_v1_20425_1_1 | GTTTACAGGGGGAGCGGTAC | TACTGCTGCTGTTGTGTTGG |
| *Cura-APC* | *Cura pinguis* | dd_Cura_v2_15083_1_1 | TCCGTCAACTTTTGTATCGACAG | TCAGCTGGAGTGTTCTCAGTC |
| *Pten-APC* | *Polycelis tenuis* | dd_Pten_v3_50650_1_2 | ACGGAAAGTCGAACAAATGCG | TCGCTGGAGTGTTCTCAGTC |
| *Ptor-APC* | *Planaria torva* | dd_Ptor_v3_37994_1_1 | TGTGAAATTCGAATCTGAACCGC | TCGTCATTAACATGGAGGGGG |
| *Smed-APC* | *Schmidtea mediterranea* | dd_Smed_v6_2688_0_1 | AACGTCTATTGTTGCCATCTTG | GCGCCAAATATCACAGTCG |
| *Smed-axin2* | *Schmidtea mediterranea* | dd_Smed_v6_5531_0_1 & dd_Smed_v6_5818_0_1 | AATCACCCCAGTTCCAGATG | TGATTTGAATTGCCCAAGC |
| *Smed-CPEB-1* | *Schmidtea mediterranea* | dd_Smes_v1_22615_1_1 | TCTTAACCGCGAGTTTAGCC | AGTGACTGCCCGCATATAAC |
| *Smed-evi* | *Schmidtea mediterranea* | dd_Smed_v6_9546_0_1 | CCATGGAGCATCGACCTCTC | ACTCCTTCGATGATGCCGTC |
| *Smed-fzd-1* | *Schmidtea mediterranea* | dd_Smed_v6_6353_0_1 | CCATTAGCAGTATGTTTCAC | ATATTCGTGCCGTTAATGAC |
| *Smed-fzd-4.2* | *Schmidtea mediterranea* | dd_Smed_v6_16926_0_1 | CAAACGTTACACATTGCACA | AAAACTCCACAAACTCTCTG |
| *Smed-fzd-4.3* | *Schmidtea mediterranea* | dd_Smed_v6_13356_0_1 | GGGCTCAGTTCACTTTTAAT | TTGCCACCAGTGAATATTTG |
| *Smed-fzd-4.4* | *Schmidtea mediterranea* | dd_Smed_v6_7210_0_1 | CTTGTATTTCTCCACGTTCA | AATGGCCATCAGAAATTTGC |
| *Smed-fzd-5/8d2* | *Schmidtea mediterranea* | dd_Smed_v6_17168_1_1 | GAAAAGACAAACCTAGACGC | CCCACCGACAATTGAAAATC |
| *Smed-fzd-5/8d4* | *Schmidtea mediterranea* | dd_Smed_v6_11823_0_1 | ATTATTATACCCTGGACACC | GTTTTCTTCTTCCCATCTAC |
| *Smed-msy-4* | *Schmidtea mediterranea* | dd_Smed_v6_25570_0_1 | GCTGCAAATGTTACGGGTCC | CATCGCTCTGGTTGCTTTCG |
| *Smed-notum* | *Schmidtea mediterranea* | dd_Smed_v6_24180_0_1 | TTAATTGAGTGAGATGTGCC | TCGACGATCACATAACTAACC |
| *Smed-ophis* | *Schmidtea mediterranea* | dd_Smed_v6_17134_0_1 | CCGCATGTAGTTGGTTAGTG | TGCAGTATTCCATGCATGGC |
| *Smed-sFRP-2* | *Schmidtea mediterranea* | dd_Smed_v6_8832_0_1 | TTTTATCGGTGTTTGTTGATCG | TTCGCCTGTTTTTATTCTGG |
| *Smed-surfactant-b* | *Schmidtea mediterranea* | dd_Smes_v1_38899_1_1 | ATCGTGCGGCAAATCTCATC | TCACCAAATCTACCGCTGCC |
| *Smed-tsp-1* | *Schmidtea mediterranea* | d_Smed_v6_69681_0_1 | CCGAAACGCACATTGTGAAATG | CTTTTGGTGGACAGCAGCTG |
| *Smed-ferritin A* | *Schmidtea mediterranea* | dd_Smes_v1_98663_1_1 | ATGACACACAGAAAAATGTCTCTCAC | TTAGAATCGATAAGTAGATCCAATTG |
| *Smed-ferritin B* | *Schmidtea mediterranea* | dd_Smes_v1_54523_1_1 | ATGTCACTTTCAAAAGTTATTGAAGTTC | AACGATTCATTGATCCAATAATAGG |
| *Smed-ferritin C* | *Schmidtea mediterranea* | dd_Smes_v1_39877_1_1 | ATGTCTTCAAAAGTTGTTGATGTTCC | CTAGAATCGAGTGGTTGATCCG |
| *Smed-ferritin D* | *Schmidtea mediterranea* | dd_Smes_v1_14451_1_1 | ACTGAAATTGAAGTCGAAGTCAGT | CTAATCTGTCGTGGATTGTTTATCA |
| *Came-ß-catenin-1* | *Camerata robusta* | dd_Came_v1_29389_1_1 | GATTTGTGCAGCCGGGATTC | CACTCCAGGTGGAATTCCCG |
| *Bcan-ß-catenin-1* | *Bdelloura candida* | dd_Bcan_v1_027080_01_01 | GCACTTGAAACCCATGCTCC | CACAGCTCACACATTCACC |
| *Cura-ß-catenin-1* | *Cura pinguis* | dd_Cura_v1_053333_01_01 | ACATGGCGGAGAAGGTATGG | ACAAACATGCACATGGGACG |
| *Pten-ß-catenin-1* | *Polycelis tenuis* | dd_Pten_v3_42913_1_1 | ATGAATGAGCAGATGAATATGGTA | CTAGGGCTTGTGGAGACGG |
| *Ptor-ß-catenin-1* | *Planaria torva* | dd_Ptor_v1_26557_0_1 | ATGAATGAACAGATGAATATCGTA | TTCAATAGTAAATCCAAGAAGTTTGAC |
| *Smed-ß-catenin-1* | *Schmidtea mediterranea* | dd_Smed_v6_2688_0_1 | CCAGATACTCCTGGTAGTAC | GACTCCAAGTGATTGAACAG |

**Supplementary Table 4.** Contig names and primers used in this study, without any vector tails (see methods for more details). **1 to 13:** to amplify specifically the F1 region of multiple species and clone it into a paff8cT4P vector for recombinant protein production. **14 to 38:** to amplify a variety of genes. **39 to 44:** to produce *ß-Catenin-1* template for *RNAi* food production. For primers **40 to 44**, the full length of the *ß-Catenin-1* gene (FL) was cloned into a pPRT4P vector. Shorter fragments were posteriorly used as a template for *RNA* production (last two primer columns).
